# Low-Density Lipoprotein Cholesterol and Cerebral Pericyte Dysfunction in Acute Ischemic Stroke: Clinical and Experimental Evidence

**DOI:** 10.64898/2026.09.27.754749

**Authors:** Feng Wang, Wenqiang Quan, Gang Wu, Jing Yang, Xu Wang, Qian Cao, Wenlin Hao, En Wang, Tomomi Furihata, Weihai Xu, Dong Li, Shaofa Ke, Yang Liu

## Abstract

**Background:** Low-density lipoprotein cholesterol (LDL-C) is an established risk factor for atherosclerotic cerebrovascular disease, but its effects on the cerebral microcirculation are poorly defined. We investigated whether LDL-C is associated with pericyte dysfunction in acute ischemic stroke (AIS) and explored underlying mechanisms.

**Methods:** Serum soluble platelet-derived growth factor receptor-β (sPDGFRβ) was measured in two independent AIS cohorts (n=97 and n=86). Cerebral pericyte transcriptional responses to LDL-C were analyzed by single-cell RNA sequencing in *Apoe*-knockout mice fed control or high-fat diet (n=4 [male/female: 2/2]/group), and cultured human brain pericytes were exposed to purified human LDL cholesterol with or without N-acetyl-L-cysteine.

**Results:** Higher serum sPDGFRβ was associated with greater neurological deficit and circulating neurofilament light chain. LDL-C positively correlated with sPDGFRβ in both clinical cohorts. High-fat diet induced coordinated suppression of type-I interferon pathways, and a positive shift in reactive oxygen species-related transcriptional programs in mouse pericytes; exploratory sex-by-diet analyses suggested larger transcriptional responses in females. In cultured human pericytes, LDL-C induced oxidative-stress, lipid-handling, and ferroptosis-associated transcriptional programs while suppressing type-I interferon signaling. Cross-species enrichment analysis showed that LDL-C-induced and -repressed signatures in cultured pericytes aligned with the sex-by-diet interaction in *Apoe*-knockout mice. LDL-C also reduced cell-surface PDGFRβ and increased sPDGFRβ release in cultured pericytes, which were significantly attenuated by N-acetyl-L-cysteine, whereas LDH activity increased numerically but not significantly.

**Conclusions:** Clinical and experimental findings support an LDL-pericyte axis linking atherogenic lipid stress to cerebral microvascular dysfunction, with possible sex-dependent modulation of oxidative stress and type-I interferon signaling as potential mechanisms.

**Clinical Perspective:** **What Is New?**

- In two independent AIS cohorts, circulating sPDGFRβ was independently associated with LDL cholesterol, linking a conventional atherosclerotic risk factor to a marker of cerebral pericyte dysfunction.
- Experimental models showed that atherogenic lipid exposure induces oxidative/lipid-stress programs and suppresses type I interferon signaling in pericytes, with a more pronounced high-fat-diet transcriptional response in females.

**What Are the Clinical Implications?**

- LDL lowering may protect the cerebral microcirculation in addition to preventing large-artery atherosclerosis, but prospective studies and stroke models are needed before pericyte-directed or sex-specific treatment implications can be established.

## Introduction

Low-density lipoprotein (LDL) cholesterol is an established risk factor for atherosclerosis and large-artery ischemic stroke, but its effects on the cerebral microcirculation are less well defined. This gap is clinically relevant because restoration of large-artery patency does not always restore tissue perfusion; persistent downstream microvascular hypoperfusion, or no-reflow, can contribute to futile recanalization and poor recovery after acute ischemic stroke (AIS) ^1–3^. In fact, there is evidence that atherogenic lipoproteins are relevant to small-vessel pathology. Small dense LDL and other ApoB-containing lipoprotein measures have been associated with white matter hyperintensities, lacunes, cerebral microbleeds, and enlarged perivascular spaces ^4, 5^. These observations raise the possibility that the vascular effects of atherogenic lipids extend beyond large arteries to the cerebral microvasculature.

Cerebral pericytes are mural cells that closely envelop capillaries and regulate capillary blood flow, microvascular architecture, and blood-brain barrier (BBB) integrity ^6^. Pericyte loss or dysfunction is associated with microvascular rarefaction and impaired tissue perfusion ^7–10^. During cerebral ischemia, pericytes contract rapidly and constrict capillaries ^11–13^. Additionally, pericyte death or detachment from the base membrane can increase BBB permeability and aggravate secondary tissue damage in ischemic brain ^14^. Hyperlipidemia has been shown to impair vascular endothelial growth factor-induced angiogenesis and reduce cerebral pericyte coverage ^15^, but whether LDL cholesterol directly alters pericyte integrity and signaling remains poorly understood.

Platelet-derived growth factor receptor-β (PDGFRβ) is highly expressed by pericytes ^16^, and its soluble ectodomain, soluble PDGFRβ (sPDGFRβ), can be detected in cerebrospinal fluid and blood after pericyte stress ^17^. Circulating sPDGFRβ and PDGFRβ-positive microvesicles have been reported after AIS ^18, 19^, supporting a dynamic pericyte response to cerebral ischemia. However, the systemic factors associated with circulating sPDGFRβ in AIS and the relationship between atherogenic lipids and pericyte dysfunction remain uncertain.

In this study, we tested the hypothesis that LDL cholesterol contributes to cerebral pericyte dysfunction in AIS. We analyzed sPDGFRβ and lipid parameters in two independent AIS cohorts, characterized cerebral pericyte transcriptional responses to high-fat feeding in *Apoe*-knockout mice, and examined the direct effects of purified human LDL cholesterol on cultured human brain pericytes. By integrating clinical, *in vivo*, and *in vitro* data, we sought to define an LDL-pericyte axis and identify molecular programs that may link dyslipidemia to cerebral microvascular vulnerability.

## Methods

### Data and materials availability statement

Deidentified patient-level data are not publicly available because of patient confidentiality and institutional regulations but may be made available by the corresponding author on reasonable request, subject to approval by the relevant ethics committees and applicable regulations. The mouse single-cell RNA-sequencing and human pericyte bulk RNA-sequencing datasets generated in this study will be deposited in the NCBI Gene Expression Omnibus after publication; linked raw sequence data are available through the Sequence Read Archive. The corresponding author had full access to all study data and takes responsibility for the integrity of the data and the accuracy of the analyses. Additional analytic code, methods, and study materials are available from the corresponding author upon reasonable request.

### Patients and study design

This prospective observational study comprised two independent cohorts of patients with AIS recruited from Taizhou Hospital and Tongji Hospital. Serum sPDGFRβ and lipid parameters were analyzed separately in each cohort; because sPDGFRβ was measured using different ELISA platforms, absolute concentrations were not compared across sites. No a priori sample-size calculation was performed because the clinical analyses were exploratory. The study protocol was approved by the Ethics Committees of Taizhou Hospital, Zhejiang Province (K20181204), and Tongji Hospital, Shanghai, China (2021-LCYJ-011). Written informed consent was obtained from patients or their legal guardians.

The Taizhou cohort included 97 patients consecutively recruited from April 2020 to September 2021, and the Tongji cohort included 86 patients consecutively recruited from January to December 2021. Inclusion criteria were age ≥18 years and a diagnosis of AIS confirmed by computed tomography or magnetic resonance imaging, consistent with the 2019 American Heart Association/American Stroke Association guideline ^20^. Acute treatment followed Chinese stroke guidelines ^21^. Exclusion criteria were non-stroke cardiovascular diseases, treatment with endovascular thrombectomy, liver or kidney disease, and infectious disease during hospitalization or within the preceding year.

### Measurement of sPDGFRβ, PDGF-BB and NfL in the serum

Whole blood was collected at 7 AM on the second day after admission and allowed to stand at room temperature for 30 to 60 minutes before centrifugation at 2000g for 15 minutes. Serum was stored at −80°C until analysis. At Taizhou Hospital, sPDGFRβ and platelet-derived growth factor-BB (PDGF-BB) were measured using ELISA kits from R&D Systems (Minneapolis, MN) and Thermo Fisher Scientific (Frederick, MD), respectively. At Tongji Hospital, sPDGFRβ and neurofilament light chain (NfL) were measured using ELISA kits from Cloud-Clone Corp. (Katy, TX). Because different sPDGFRβ assays were used at the two sites, associations were evaluated within each cohort rather than by directly comparing absolute concentrations across cohorts.

### Clinical data collection

Demographic characteristics, medical history, clinical examination findings, laboratory results, and diagnoses were extracted from electronic medical records. All patients in the Taizhou cohort completed 3-month follow-up, with no loss to follow-up. Functional outcome at 90 days was assessed with the modified Rankin Scale (mRS); favorable outcome was defined as mRS <2. Neurological deficit was assessed with the National Institutes of Health Stroke Scale (NIHSS) at admission and after 7 days or at hospital discharge.

### Mice and diet

Twenty-one 2-month-old *Apoe*-knockout (*Apoe*^-/-^; B6.129P2-Apoetm1Unc/J) ^22^ littermate mice, back-crossed to a C57BL/6J genetic background for more than 6 generations, were purchased from Hangzhou Hans Biotechnology (Hangzhou, China). Mice were allocated to standard chow (CTRL) or a high-fat diet (HFD) for 2 months within each sex while preserving littermate matching between dietary groups, so that diet effects were compared between animals of the same sex and comparable genetic background. No computer-generated randomization sequence was used. Animals were maintained under standard institutional animal-housing conditions with a conventional light–dark cycle; no additional study-specific environmental enrichment was introduced beyond routine husbandry. Animals were monitored throughout the feeding period as part of routine animal care for general health and signs of distress, and humane endpoints and veterinary intervention followed the approved institutional animal-welfare protocol. Blinding during animal handling and diet administration was not feasible because high-fat feeding produced readily visible phenotypic differences, including increased body size. For single-cell RNA sequencing, two mice per sex×diet group were selected from the larger cohort while preserving littermate matching; selection was not based on serum cholesterol concentration or other measured outcomes.

The HFD was based on the D12492 formulation (Hangzhou Hans Biotechnology) and provided 60% of total energy from fat (predominantly lard with a smaller proportion of soybean oil), 20% from carbohydrate, and 20% from protein. Diet-induced hypercholesterolemia was validated by measuring serum total and LDL cholesterol in all 21 mice as a commercial service at Shanghai Model Organisms (Shanghai, China) (Supplementary Figure S1), and the atherogenic phenotype was further confirmed by Oil Red O staining of the aortic arch (Supplementary Figure S2).

Animal procedures complied with national regulations and ARRIVE guidelines and were approved by the Ethics Committee for Animal Welfare at Taizhou Hospital, Zhejiang Province (registration number: tzy-2024006).

### Brain tissue collection for single-cell sequencing

Mice were euthanized by isoflurane overdose and perfused with ice-cold Dulbecco’s phosphate-buffered saline (DPBS). Cortical and hippocampal tissue from 8 littermate mice was carefully dissected and transported on ice in the MACS® Tissue Storage Solution (Miltenyi Biotec China, Shanghai, China; Catalog number: 130-100-008) to Majorbio Bio-pharm Technology Co. Ltd (Shanghai, China) for single-cell processing.

### Tissue dissociation and cell purification

Brain tissue was minced on ice and dissociated with 0.25% trypsin (Thermo Fisher Scientific; Catalog number: 25200-072) and 10 μg/mL DNase I (Sigma-Aldrich; Catalog number: 11284932001) in DPBS containing 5% fetal bovine serum at 37°C with gentle shaking for approximately 40 minutes. Cell suspensions were passed through a 40-μm nylon strainer, and red blood cells were removed with 1× Red Blood Cell Lysis Solution (Thermo Fisher Scientific; Catalog number: 00-4333-57). After washing in DPBS containing 0.4% fetal bovine serum, cell viability was assessed with AO/PI staining. Single-cell libraries were prepared using the DNBelab C TaiM4 V3.0 workflow (MGI Tech Co., Ltd., Shenzhen, China) according to the manufacturer’s protocol and sequenced on the DNBseq platform using 2×100-bp chemistry.

### Single-cell RNA sequencing analysis

Reads were processed with the dnbc4tools v2.1.2 pipeline using default and recommended parameters and aligned to the GRCm39 mouse reference genome with STAR ^23^.

**1) Preprocessing and quality control.** Background-corrected count matrices (barcodes, features, and matrix triplets; GRCm39, Ensembl) from eight animals were imported into R (version 4.6.1) using Seurat (version 5.5.1). Genes detected in fewer than 3 cells were removed. Cells were retained if they expressed ≥500 genes and contained <10% mitochondrial and <5% hemoglobin transcripts; the input matrices had already been filtered to contain at least 1000 UMIs per barcode. Doublets were identified and removed for each sample using scDblFinder (version 1.26.7) with default parameters.

**2) Integration, clustering, and annotation.** Filtered objects were merged and log-normalized, and 2000 variable features were selected. After scaling and principal component analysis using 30 principal components, batch effects between animals were corrected with Harmony (version 2.0.5; group.by.vars = “sample”). Sex and diet were not used as integration covariates to preserve biological contrasts. A shared-nearest-neighbor graph and Louvain clustering (resolution 0.6) were used, and UMAP was computed on the Harmony embedding. Clusters were annotated with module scores based on curated marker panels for pericytes, smooth muscle cells, endothelial cells, astrocytes, microglia, neurons, oligodendrocytes, oligodendrocyte precursor cells, fibroblasts, ependymal cells, and T cells, followed by verification with canonical marker expression.

**3) Pericyte refinement and decontamination.** Cells in clusters annotated as pericytes were reclustered (resolution 1.2). Subclusters dominated by non-mural cell signatures were removed, and remaining cells were retained only when their highest-scoring signature corresponded to pericytes or smooth muscle cells. Ambient RNA was estimated and subtracted with DecontX (celda version 1.28.0) using cell-type labels as z and animal as batch. Cells with estimated contamination > 0.5 were excluded. Residual contamination was quantified from canonical microglial and endothelial marker sets relative to the corresponding source-cell populations.

**4) Differential expression**. Decontaminated counts were summed within each animal to generate pseudobulk profiles, rounded to whole numbers, and genes with ≥ 10 counts in ≥ 4 samples were retained. DESeq2 (version 1.52.0) was run with an additive design (∼ sex + diet) to estimate sex-adjusted diet and diet-adjusted sex effects and with an interaction design (∼ sex × diet) to evaluate sex-dependent diet responses. Multiple comparisons were controlled with the Benjamini-Hochberg procedure.

**5) Gene set enrichment**. Genes were ranked by the DESeq2 Wald statistic after exclusion of ribosomal genes (^Rp[sl], ^Mrp[sl]). Gene-set enrichment analysis (GSEA) was performed with clusterProfiler (version 4.20.0) using the MSigDB Mouse Hallmark and Reactome collections (msigdbr version 26.1.1; minGSSize=15, maxGSSize=500) and a curated type I interferon module. To test robustness to residual contamination, pseudobulk aggregation, DESeq2 modeling, and GSEA were repeated in the 50% of pericytes with the lowest DecontX contamination estimates.

**6) Cross-dataset analysis.** Genes differentially expressed in LDL-treated versus control human pericytes (FDR <0.05 and |log2 fold change| ≥1) were separated into LDL-induced and LDL-repressed signatures (see the following sections on cultured human pericytes). Human genes were mapped to one-to-one mouse orthologs using babelgene (version 22.9). Each signature was tested by GSEA against the *in vivo* sex-adjusted diet ranking and the sex-by-diet interaction ranking. Gene-level concordance was assessed by Spearman correlation between the *in vitro* log2 fold change and the corresponding *in vivo* Wald statistic across shared orthologs.

**7) Statistics and reproducibility**. The mouse experiment included 2 animals per sex-by-diet group (8 animals total), and the transcriptomic pericyte culture experiment included 6 independent samples per group (12 independent experiments total). Pseudobulk aggregation was used so that the animal, rather than the individual cell, was the unit of biological replication. An adjusted *p* value <0.05 was considered significant. Given the small group size, sex-by-diet interaction results are interpreted at the level of coordinated gene sets and considered hypothesis-generating.

### Human brain pericyte culture and treatments

A conditionally immortalized human brain vascular pericyte line (HBPC, clone ci37) was cultured at 33°C in 5% CO2/95% air in pericyte medium (ScienCell Research Laboratories; Catalog number: 1201) containing 2% fetal bovine serum, 1% pericyte growth factors, and penicillin-streptomycin, as described previously ^24^. Culture flasks and plates were coated with rat tail collagen type I (Thermo Fisher Scientific; Catalog number: A1048301). For LDL exposure experiments, pericytes were cultured in 24-well plate at 2.0 × 10^5^ cells/well. Before experiments, the culture medium was replaced with serum-free pericyte medium and cells were cultured at 37°C for 3 days to facilitate the cell differentiation ^25^. Pericytes were then treated with purified LDL from human plasma (Sigma-Aldrich; Catalog number: SAE0053) at 0, 1 and 2 mg/ml for 24 hours, with or without 2 mmol/L N-acetyl-L-cysteine (Sigma-Aldrich; Catalog number: A9165) using our established protocol ^26^. Cell-surface PDGFRβ was quantified by flow cytometry using a PE-conjugated anti-human PDGFRβ antibody (Miltenyi Biotec; Clone: REA363). Conditioned medium was collected for sPDGFRβ ELISA (R&D Systems) and lactate dehydrogenase (LDH) assays using CyQUANT™ LDH Cytotoxicity Assays (Thermo Fisher Scientific). Because the LDL preparation could contribute background LDH activity, cell-free medium containing 1 or 2 mg/mL LDL was assayed in parallel, and these values were subtracted from the corresponding conditioned-medium measurements. After removal of the culture medium, cells were immediately lysed in RLT Plus buffer from the RNeasy Plus Mini Kit (Qiagen, Hilden, Germany) for RNA isolation.

### mRNA sequencing and transcriptome analysis

Total RNA was isolated from cultured pericytes using the RNeasy Plus Mini Kit (Qiagen). Poly(A)-enriched RNA-sequencing libraries were generated and sequenced on an Illumina platform to a target depth of approximately 20 million paired-end 2×150-bp reads per sample. After adapter and quality filtering, reads were aligned to the human GRCh38 reference genome with HISAT2 ^27^ and assembled with StringTie ^28^. Differential expression was analyzed with DESeq2; differentially expressed genes were defined as |log2 fold change| ≥1 and FDR <0.01. Gene Ontology and Kyoto Encyclopedia of Genes and Genomes enrichment analyses were then performed. Library preparation, sequencing, and primary transcriptomic analysis were conducted by Biomarker Technologies (BMK) GmbH (Münster, Germany).

### Quantitative PCR for analysis of gene transcripts

Total RNA was reverse-transcribed. Gene transcripts were quantified with established protocols^29^. The transcription of *AIFM2*, *AKR1C1*, *AKR1C2*, *DDIT3*, *FTH1*, *FTL*, *GCLC*, *GCLM*, *IFI35*, *NQO1*, *PCSK9*, *PLIN2*, *SAMHD1*, *SLC7A11*, *STAT1*, and *SRXN1* was determined to validate the results from mRNA sequencing using the SYBR green binding technique with the primers listed in Supplementary Table 1. TaqMan gene expression assays of 18S ribosomal RNA (*18S*) (Thermo Fisher Scientific) was used as an internal control.

### Statistical analysis

Statistical analyses were performed using SPSS for Windows (version 26.0; IBM, Armonk, NY, USA). Continuous variables are presented as mean ± SD for clinical data and mean ± SEM for experimental data, whereas categorical variables are presented as frequencies and percentages. Continuous variables were compared between two independent groups using Student’s *t*-test or the Mann–Whitney *U* test, as appropriate. Categorical variables were compared using Pearson’s χ² test. All tests were two-sided, and *p* < 0.05 was considered statistically significant.

To examine the association between serum sPDGFRβ and neurological severity in the Taizhou cohort, patients were categorized according to NIHSS score (< 4 vs ≥ 4), which resulted in two groups with nearly the same patient number per group. Candidate clinical and laboratory variables were first evaluated in univariable analyses. Variables associated with the NIHSS category at *p* < 0.10 were considered candidates for multivariable logistic regression to determine whether serum sPDGFRβ was independently associated with neurological severity.

Within each AIS cohort, associations between serum sPDGFRβ and continuous clinical or laboratory variables were evaluated using Pearson’s or Spearman’s correlation analysis, as appropriate according to data distribution. For categorical variables, serum sPDGFRβ concentrations were compared using Student’s *t*-test or analysis of variance for normally distributed data and the Mann–Whitney *U* test or Kruskal–Wallis test for non-normally distributed data, as appropriate. Variables associated with serum sPDGFRβ at *p* < 0.10 in univariable analyses were considered candidate variables for multivariable linear regression.

Candidate variables were initially entered simultaneously into a full multivariable linear regression model. Multicollinearity was assessed using tolerance, variance inflation factor (VIF), and collinearity diagnostics before interpretation of the regression coefficients. Variables representing strongly overlapping biological or mathematical measures were not retained simultaneously when substantial multicollinearity was identified. Specifically, total cholesterol, high-density lipoprotein (HDL) cholesterol and triglyceride were not included together with LDL cholesterol when marked collinearity between these lipid measures was observed. Similarly, simultaneous inclusion of total white blood cell (WBC) count and its differential leukocyte components resulted in severe multicollinearity; therefore, differential leukocyte counts were excluded from the revised model and total WBC count was retained as the overall measure of circulating leukocytes. The revised full model was refitted after resolution of multicollinearity. A backward elimination procedure, using *p* ≥ 0.10 as the criterion for removal, was subsequently applied to the revised candidate-variable set to derive a parsimonious model and assess the stability of the observed associations. Unstandardized regression coefficients (B), standardized β coefficients, 95% confidence intervals (CIs), and *p* values are reported for multivariable linear regression analyses.

In hypercholesterolemia mice, total and LDL cholesterol were compared between dietary groups within each sex using a two-way ANOVA, with diet and sex as fixed factors, followed by a Tukey test for multiple comparisons between all four group means (Supplementary Fig. S1, A, B); the homogeneity of variances was confirmed by Levene’s test. To compare the magnitude of the diet response between the two cholesterol fractions, which differ approximately five-fold in absolute concentration, concentrations were log₂-transformed so that the diet effect is expressed as a fold-change. Fold-changes with 95% confidence intervals (Supplementary Fig. S1, C) were estimated from linear models on log₂ concentrations adjusted for sex. Because both fractions were measured in the same animal, the two outcomes were analyzed jointly in a linear mixed-effects model (analyte × diet + analyte × sex, random intercept per animal, REML); on the log scale the analyte × diet interaction equals the ratio of the two-fold-changes and therefore tests directly whether the relative LDL cholesterol response exceeds that of total cholesterol. This result was confirmed by exact permutation and by leave-one-animal-out refitting. The analyses were performed using GraphPad Prism Version 8.0.2 and Python 3.11.

In human pericyte culture, the independent experiment was considered the unit of biological replication; technical replicates, where present, were averaged within each experiment before statistical analysis. For flow-cytometric analysis of cell-surface PDGFRβ, four independent experiments were performed, with 2 mg/mL LDL condition unavailable in one experiment; therefore, the factorial analysis was performed in the three experiments with complete measurements at all LDL concentrations. LDL concentration (0, 1, and 2 mg/mL) and NAC treatment (absent or present) were treated as within-experiment factors and analyzed using two-way repeated-measures ANOVA. Greenhouse–Geisser correction was applied to effects involving the three-level LDL factor. The LDL×NAC interaction was used to test whether NAC modified the LDL concentration-response relationship.

Extracellular sPDGFRβ and LDH activity were measured in five independent experiments with complete measurements for all LDL and NAC conditions and were analyzed using the same 2×3 repeated-measures design. When an LDL effect was examined within an individual NAC condition, one-way repeated-measures ANOVA with Greenhouse–Geisser correction was used, followed where appropriate by paired comparisons with Holm adjustment for multiple testing. Because sPDGFRβ concentrations were strictly positive and showed substantial between-experiment variation in baseline magnitude, a log-transformed sensitivity analysis was additionally performed to evaluate proportional treatment effects. For the 2 mg/mL LDL condition, proportional responses were also expressed as the LDL2/LDL0 fold change within each experiment and NAC condition, and the ratio of these fold changes was evaluated on the logarithmic scale. Effect sizes are reported as partial η². All tests were conducted with R (version 4.6.1).

## Results

### Serum sPDGFRβ was associated with neurological severity and NfL in AIS

We first examined whether serum sPDGFRβ was associated with neurological severity in the Taizhou cohort (n = 97). Patients with NIHSS ≥ 4 had higher serum sPDGFRβ concentrations than those with NIHSS < 4 (Fig. 1, A; *t*-test, *p* < 0.05). Age, white blood cell and neutrophil counts, D-dimer, and total bilirubin also differed between NIHSS groups (Supplementary Table 2; *t*-test or Mann-Whitney *U* test, *p* < 0.05). In a multivariable logistic regression model including candidate variables with *p* < 0.10 in univariable analyses, serum sPDGFRβ remained independently associated with the NIHSS category (Supplementary Table 3; Logistic regression, *p* < 0.05).

**Figure 1.**
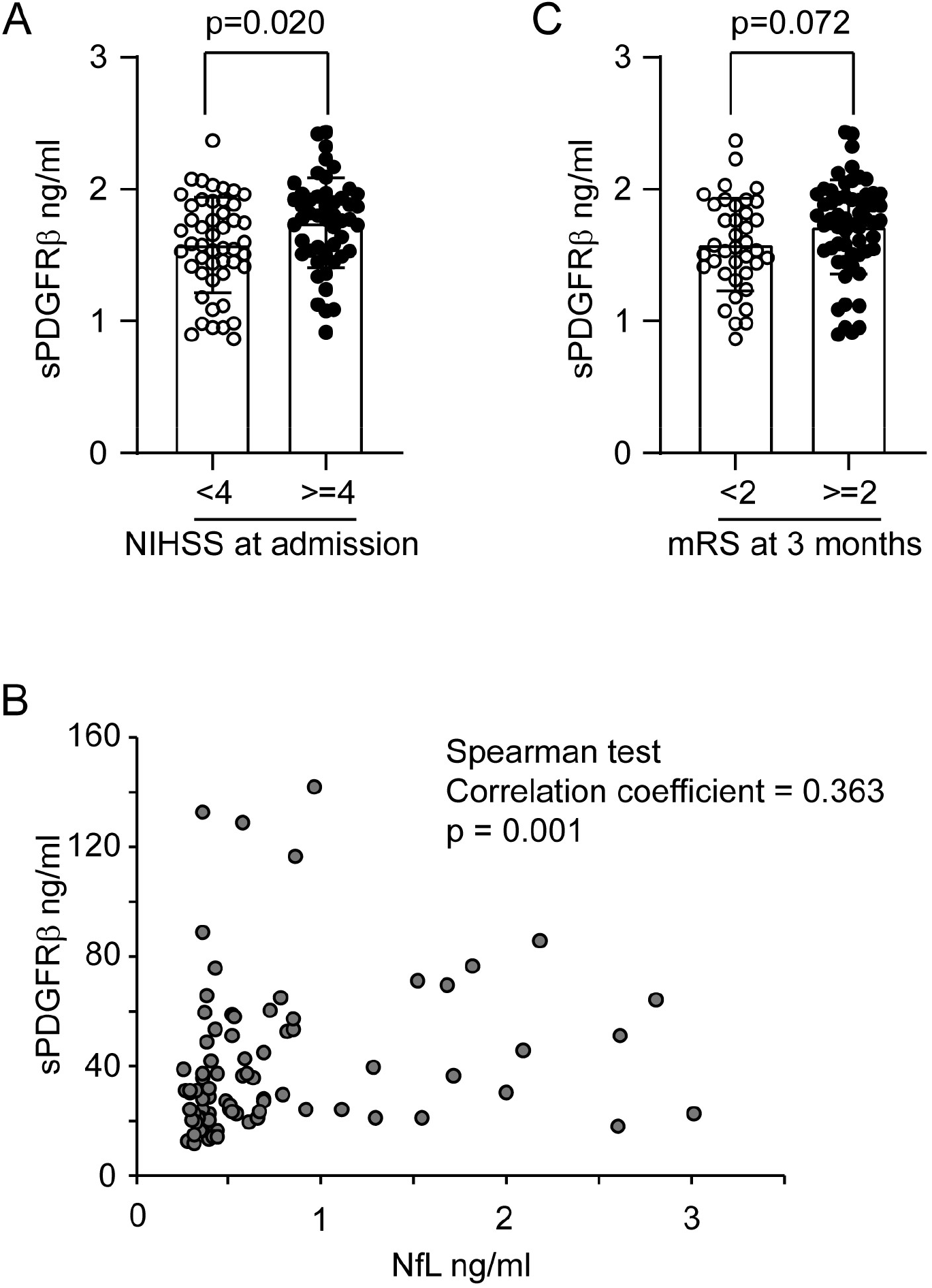
Serum sPDGFRβ in relation to neurological severity, NfL, and 3-month outcome after AIS. Serum sPDGFRβ was analyzed in patients with AIS from Taizhou Hospital (A and C) and Tongji Hospital (B). sPDGFRβ concentrations were higher in patients with NIHSS ≥4 than in those with NIHSS <4 in the Taizhou cohort (A; n=49 and 48, respectively; independent-samples *t* test). In the Tongji cohort, serum sPDGFRβ correlated positively with neurofilament light chain (NfL) (B; n=86; Spearman correlation). In the Taizhou cohort, sPDGFRβ tended to be lower in patients with favorable 3-month outcome (mRS <2) than in those with mRS ≥2 (C; n=38 and 59, respectively; independent-samples *t* test). AIS indicates acute ischemic stroke; mRS, modified Rankin Scale; NIHSS, National Institutes of Health stroke scale; and sPDGFRβ, soluble platelet-derived growth factor receptor-β.

In the Tongji cohort (n = 86), serum sPDGFRβ was positively correlated with serum NfL (Fig. 1, B; Spearman correlation, *p* < 0.05), supporting an association between the circulating pericyte-related marker and axonal injury ^30^.

In the Taizhou cohort, patients with favorable 3-month functional outcome (mRS <2) tended to have lower serum sPDGFRβ concentrations than patients with mRS ≥2, although the difference did not reach statistical significance (Fig. 1, C; *p* = 0.072). Because this outcome analysis was exploratory, no adjusted prognostic model was performed.

### Serum sPDGFRβ was independently associated with LDL cholesterol in two AIS cohorts

We next examined clinical and metabolic factors associated with serum sPDGFRβ. In the Taizhou cohort, sPDGFRβ correlated in univariable analyses with age, alcohol use, glucose, and LDL cholesterol (Table 1, Pearson correlation/one-way ANOVA; *p* < 0.05), and showed a trend with total cholesterol (*p* = 0.077).

**Table 1.**
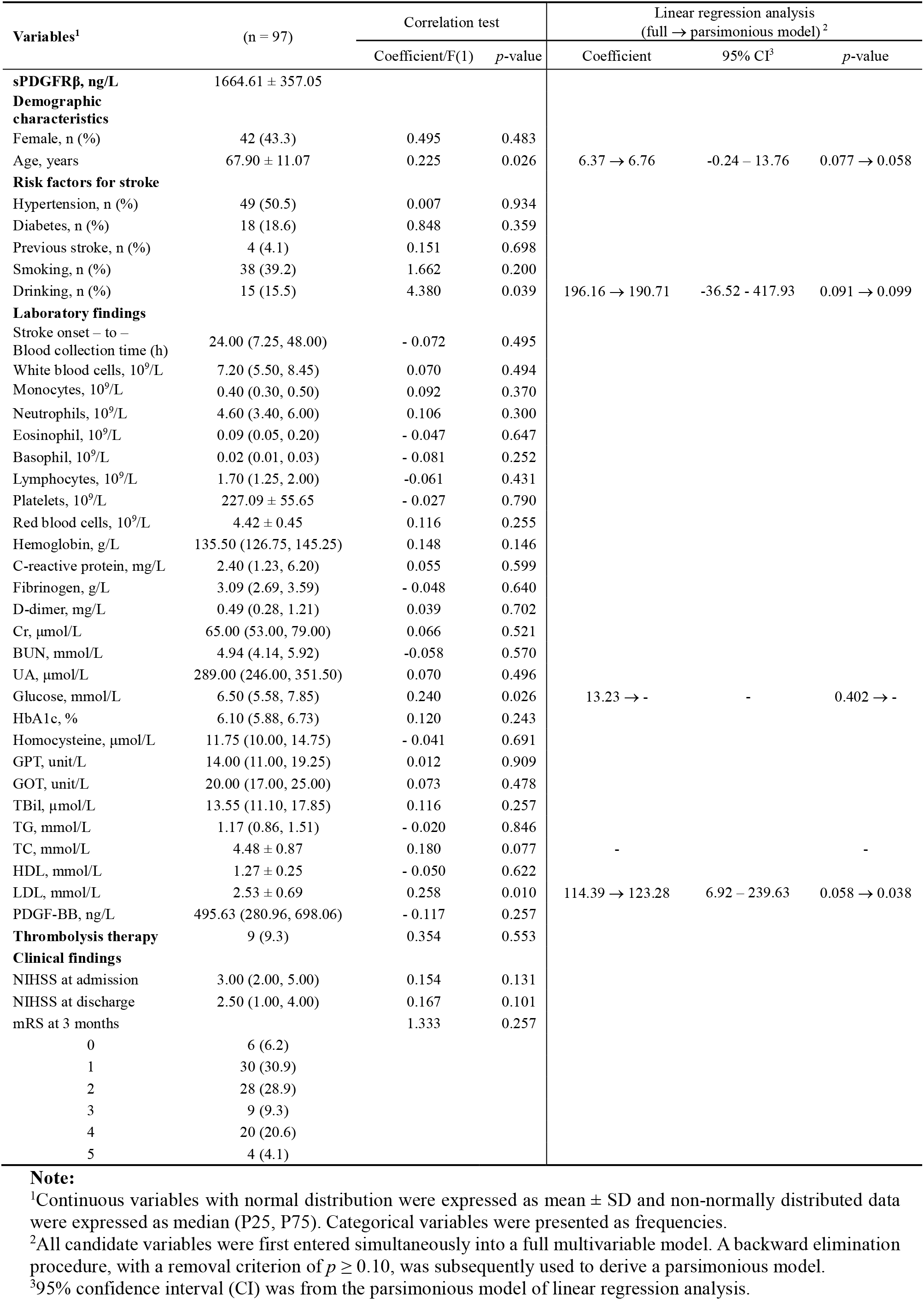
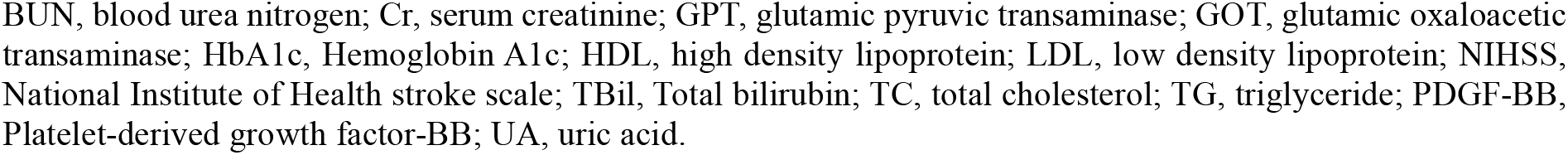
Clinical characteristics and associations with serum sPDGFRβ in the Taizhou cohort.

| Variables<sup>1</sup> | (n = 97) | Correlation test | | Linear regression analysis<br>(full → parsimonious model)<sup>2</sup> | | |
| --- | --- | --- | --- | --- | --- | --- |
|  |  | Coefficient/F(1) | p-value | Coefficient | 95% CI<sup>3</sup> | p-value |
| **sPDGFRβ, ng/L** | 1664.61 ± 357.05 | | | | | |
| **Demographic characteristics** |  |  |  |  |  |  |
| Female, n (%) | 42 (43.3) | 0.495 | 0.483 |  |  |  |
| Age, years | 67.90 ± 11.07 | 0.225 | 0.026 | 6.37 → 6.76 | -0.24 – 13.76 | 0.077 → 0.058 |
| **Risk factors for stroke** |  |  |  |  |  |  |
| Hypertension, n (%) | 49 (50.5) | 0.007 | 0.934 |  |  |  |
| Diabetes, n (%) | 18 (18.6) | 0.848 | 0.359 |  |  |  |
| Previous stroke, n (%) | 4 (4.1) | 0.151 | 0.698 |  |  |  |
| Smoking, n (%) | 38 (39.2) | 1.662 | 0.200 |  |  |  |
| Drinking, n (%) | 15 (15.5) | 4.380 | 0.039 | 196.16 → 190.71 | -36.52 – 417.93 | 0.091 → 0.099 |
| **Laboratory findings** |  |  |  |  |  |  |
| Stroke onset – to –<br>Blood collection time (h) | 24.00 (7.25, 48.00) | - 0.072 | 0.495 |  |  |  |
| White blood cells, 10<sup>9</sup>/L | 7.20 (5.50, 8.45) | 0.070 | 0.494 |  |  |  |
| Monocytes, 10<sup>9</sup>/L | 0.40 (0.30, 0.50) | 0.092 | 0.370 |  |  |  |
| Neutrophils, 10<sup>9</sup>/L | 4.60 (3.40, 6.00) | 0.106 | 0.300 |  |  |  |
| Eosinophil, 10<sup>9</sup>/L | 0.09 (0.05, 0.20) | - 0.047 | 0.647 |  |  |  |
| Basophil, 10<sup>9</sup>/L | 0.02 (0.01, 0.03) | - 0.081 | 0.252 |  |  |  |
| Lymphocytes, 10<sup>9</sup>/L | 1.70 (1.25, 2.00) | -0.061 | 0.431 |  |  |  |
| Platelets, 10<sup>9</sup>/L | 227.09 ± 55.65 | - 0.027 | 0.790 | | | |
| Red blood cells, 10<sup>9</sup>/L | 4.42 ± 0.45 | 0.116 | 0.255 | | | |
| Hemoglobin, g/L | 135.50 (126.75, 145.25) | 0.148 | 0.146 |  |  |  |
| C-reactive protein, mg/L | 2.40 (1.23, 6.20) | 0.055 | 0.599 |  |  |  |
| Fibrinogen, g/L | 3.09 (2.69, 3.59) | - 0.048 | 0.640 |  |  |  |
| D-dimer, mg/L | 0.49 (0.28, 1.21) | 0.039 | 0.702 |  |  |  |
| Cr, μmol/L | 65.00 (53.00, 79.00) | 0.066 | 0.521 | | | |
| BUN, mmol/L | 4.94 (4.14, 5.92) | -0.058 | 0.570 |  |  |  |
| UA, μmol/L | 289.00 (246.00, 351.50) | 0.070 | 0.496 | | | |
| Glucose, mmol/L | 6.50 (5.58, 7.85) | 0.240 | 0.026 | 13.23 → - | - | 0.402 → - |
| HbA1c, % | 6.10 (5.88, 6.73) | 0.120 | 0.243 |  |  |  |
| Homocysteine, μmol/L | 11.75 (10.00, 14.75) | - 0.041 | 0.691 | | | |
| GPT, unit/L | 14.00 (11.00, 19.25) | 0.012 | 0.909 |  |  |  |
| GOT, unit/L | 20.00 (17.00, 25.00) | 0.073 | 0.478 |  |  |  |
| TBil, μmol/L | 13.55 (11.10, 17.85) | 0.116 | 0.257 | | | |
| TG, mmol/L | 1.17 (0.86, 1.51) | - 0.020 | 0.846 |  |  |  |
| TC, mmol/L | 4.48 ± 0.87 | 0.180 | 0.077 | - | | - |
| HDL, mmol/L | 1.27 ± 0.25 | - 0.050 | 0.622 | | | |
| LDL, mmol/L | 2.53 ± 0.69 | 0.258 | 0.010 | 114.39 → 123.28 | 6.92 – 239.63 | 0.058 → 0.038 |
| PDGF-BB, ng/L | 495.63 (280.96, 698.06) | - 0.117 | 0.257 |  |  |  |
| **Thrombolysis therapy** | 9 (9.3) | 0.354 | 0.553 |  |  |  |
| **Clinical findings** |  |  |  |  |  |  |
| NIHSS at admission | 3.00 (2.00, 5.00) | 0.154 | 0.131 |  |  |  |
| NIHSS at discharge | 2.50 (1.00, 4.00) | 0.167 | 0.101 |  |  |  |
| mRS at 3 months |  | 1.333 | 0.257 |  |  |  |
| 0 | 6 (6.2) |  |  |  |  |  |
| 1 | 30 (30.9) |  |  |  |  |  |
| 2 | 28 (28.9) |  |  |  |  |  |
| 3 | 9 (9.3) |  |  |  |  |  |
| 4 | 20 (20.6) |  |  |  |  |  |
| 5 | 4 (4.1) |  |  |  |  |  |

**Note:**
<sup>1</sup>Continuous variables with normal distribution were expressed as mean ± SD and non-normally distributed data were expressed as median (P25, P75). Categorical variables were presented as frequencies.
<sup>2</sup>All candidate variables were first entered simultaneously into a full multivariable model. A backward elimination procedure, with a removal criterion of $p \geq 0.10$ , was subsequently used to derive a parsimonious model.
<sup>3</sup>95% confidence interval (CI) was from the parsimonious model of linear regression analysis.
BUN, blood urea nitrogen; Cr, serum creatinine; GPT, glutamic pyruvic transaminase; GOT, glutamic oxaloacetic transaminase; HbA1c, Hemoglobin A1c; HDL, high density lipoprotein; LDL, low density lipoprotein; NIHSS, National Institute of Health stroke scale; TBil, Total bilirubin; TC, total cholesterol; TG, triglyceride; PDGF-BB, Platelet-derived growth factor-BB; UA, uric acid.

In the full multivariable model including age, alcohol consumption, glucose, and LDL cholesterol as independent variables, higher LDL cholesterol showed a positive association with serum sPDGFRβ levels, although the association narrowly missed conventional statistical significance (B = 114.4, standardized β = 0.202, 95% CI −4.1 to 232.8, *p* = 0.058). Total cholesterol was not entered the multivariable analysis due to its sufficient overlap with LDL cholesterol. The full multivariable model showed that glucose did not contribute to the model (*p* = 0.402) and was removed during backward elimination in the following parsimonious model. In the parsimonious model including age, alcohol consumption, and LDL cholesterol, LDL cholesterol was the only remained variable independently associated with higher serum sPDGFRβ levels (B = 123.3, standardized β = 0.218, 95% CI 6.9 to 239.6, *p* = 0.038). The LDL effect estimate was similar between the full and parsimonious models, and no evidence of multicollinearity was observed (VIF for LDL cholesterol = 1.001 in the final model). (Table 1, Linear regression; both *p* < 0.05).

Since pericyte activation may evolve during acute ischemia ^19^, we also examined whether variation in the interval from stroke onset to blood sampling affected sPDGFRβ. No significant association was observed between sPDGFRβ and the stroke-onset-to-blood-collection interval in the Taizhou cohort (Table 1; Pearson correlation; *p* > 0.05).

The association of serum sPDGFRβ with LDL cholesterol was reproduced in the independent Tongji cohort. In univariable analyses, serum sPDGFRβ correlated with several hematologic and metabolic variables, including white blood cells (WBCs), monocytes, neutrophils, lymphocytes, platelets, red blood cells, creatinine, triglycerides, LDL cholesterol, and NfL (Table 2, Pearson correlation; *p* < 0.05).

**Table 2.**
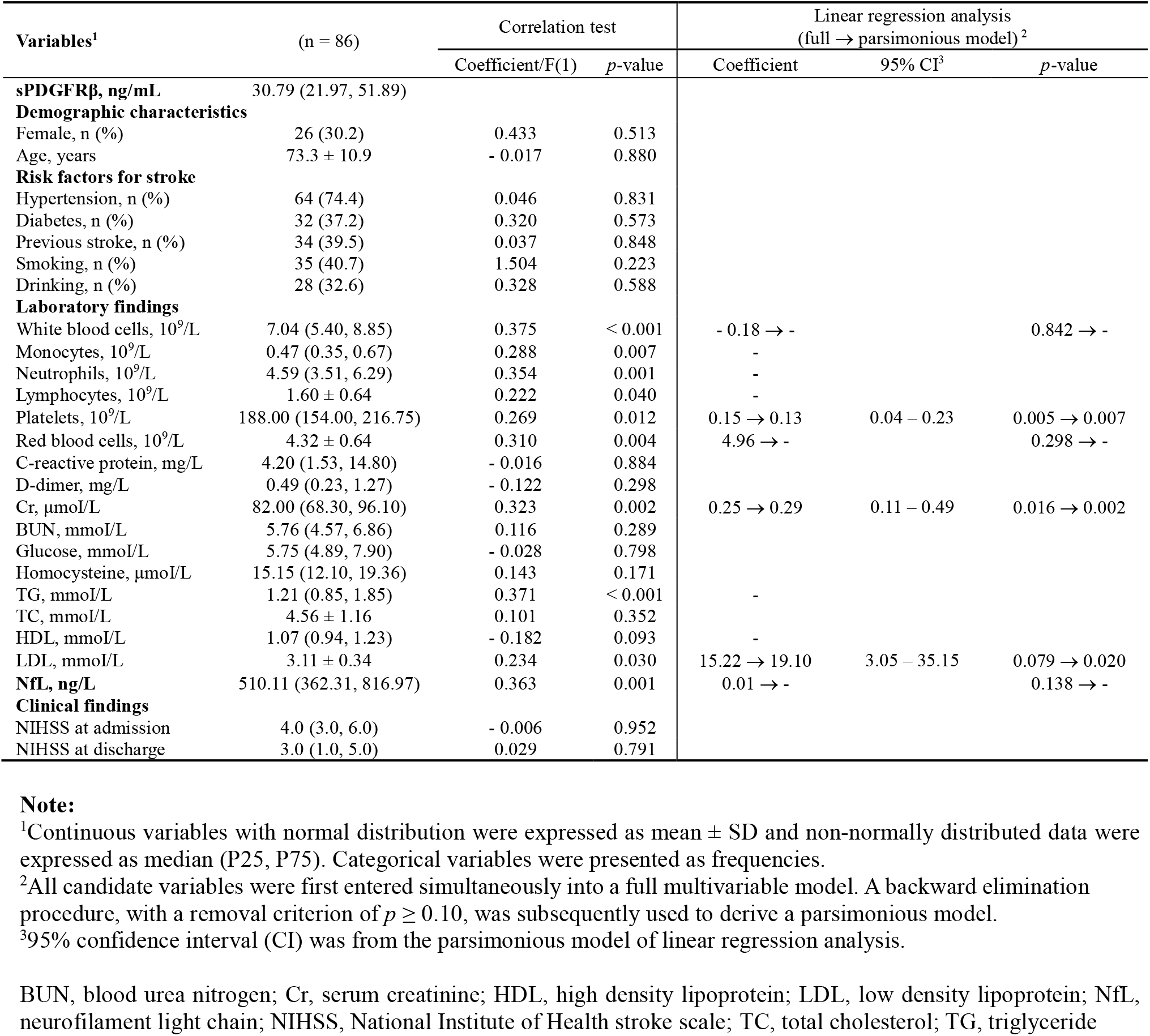
Clinical characteristics and associations with serum sPDGFRβ in the Tongji cohort.

| Variables<sup>1</sup> | (n = 86) | Correlation test | | Linear regression analysis<br>(full → parsimonious model)<sup>2</sup> | | |
| --- | --- | --- | --- | --- | --- | --- |
|  |  | Coefficient/F(1) | p-value | Coefficient | 95% CI<sup>3</sup> | p-value |
| **sPDGFRβ, ng/mL** | 30.79 (21.97, 51.89) |  |  |  |  |  |
| **Demographic characteristics** |  |  |  |  |  |  |
| Female, n (%) | 26 (30.2) | 0.433 | 0.513 |  |  |  |
| Age, years | 73.3 ± 10.9 | - 0.017 | 0.880 | | | |
| **Risk factors for stroke** |  |  |  |  |  |  |
| Hypertension, n (%) | 64 (74.4) | 0.046 | 0.831 |  |  |  |
| Diabetes, n (%) | 32 (37.2) | 0.320 | 0.573 |  |  |  |
| Previous stroke, n (%) | 34 (39.5) | 0.037 | 0.848 |  |  |  |
| Smoking, n (%) | 35 (40.7) | 1.504 | 0.223 |  |  |  |
| Drinking, n (%) | 28 (32.6) | 0.328 | 0.588 |  |  |  |
| **Laboratory findings** |  |  |  |  |  |  |
| White blood cells, 10<sup>9</sup>/L | 7.04 (5.40, 8.85) | 0.375 | < 0.001 | - 0.18 → - | | 0.842 → - |
| Monocytes, 10<sup>9</sup>/L | 0.47 (0.35, 0.67) | 0.288 | 0.007 | - |  |  |
| Neutrophils, 10<sup>9</sup>/L | 4.59 (3.51, 6.29) | 0.354 | 0.001 | - |  |  |
| Lymphocytes, 10<sup>9</sup>/L | 1.60 ± 0.64 | 0.222 | 0.040 | - | | |
| Platelets, 10<sup>9</sup>/L | 188.00 (154.00, 216.75) | 0.269 | 0.012 | 0.15 → 0.13 | 0.04 – 0.23 | 0.005 → 0.007 |
| Red blood cells, 10<sup>9</sup>/L | 4.32 ± 0.64 | 0.310 | 0.004 | 4.96 → - | | 0.298 → - |
| C-reactive protein, mg/L | 4.20 (1.53, 14.80) | - 0.016 | 0.884 |  |  |  |
| D-dimer, mg/L | 0.49 (0.23, 1.27) | - 0.122 | 0.298 |  |  |  |
| Cr, μmol/L | 82.00 (68.30, 96.10) | 0.323 | 0.002 | 0.25 → 0.29 | 0.11 – 0.49 | 0.016 → 0.002 |
| BUN, mmol/L | 5.76 (4.57, 6.86) | 0.116 | 0.289 |  |  |  |
| Glucose, mmol/L | 5.75 (4.89, 7.90) | - 0.028 | 0.798 |  |  |  |
| Homocysteine, μmol/L | 15.15 (12.10, 19.36) | 0.143 | 0.171 | | | |
| TG, mmol/L | 1.21 (0.85, 1.85) | 0.371 | < 0.001 | - |  |  |
| TC, mmol/L | 4.56 ± 1.16 | 0.101 | 0.352 | | | |
| HDL, mmol/L | 1.07 (0.94, 1.23) | - 0.182 | 0.093 | - |  |  |
| LDL, mmol/L | 3.11 ± 0.34 | 0.234 | 0.030 | 15.22 → 19.10 | 3.05 – 35.15 | 0.079 → 0.020 |
| **NfL, ng/L** | 510.11 (362.31, 816.97) | 0.363 | 0.001 | 0.01 → - | | 0.138 → - |
| **Clinical findings** |  |  |  |  |  |  |
| NIHSS at admission | 4.0 (3.0, 6.0) | - 0.006 | 0.952 |  |  |  |
| NIHSS at discharge | 3.0 (1.0, 5.0) | 0.029 | 0.791 |  |  |  |

**Note:**
<sup>1</sup>Continuous variables with normal distribution were expressed as mean ± SD and non-normally distributed data were expressed as median (P25, P75). Categorical variables were presented as frequencies.
<sup>2</sup>All candidate variables were first entered simultaneously into a full multivariable model. A backward elimination procedure, with a removal criterion of $p \geq 0.10$ , was subsequently used to derive a parsimonious model.
<sup>3</sup>95% confidence interval (CI) was from the parsimonious model of linear regression analysis.
BUN, blood urea nitrogen; Cr, serum creatinine; HDL, high density lipoprotein; LDL, low density lipoprotein; NfL, neurofilament light chain; NIHSS, National Institute of Health stroke scale; TC, total cholesterol; TG, triglyceride

In initial multivariable regression analysis, when total WBC and differential leukocyte counts were entered simultaneously, severe multicollinearity was observed. The regression model was therefore revised to retain total WBC count while excluding its differential leukocyte components. As triglyceride was closely correlated with LDL cholesterol (Pearson correlation, *p* < 0.001), it was also excluded for multivariable analysis. In the revised full model including WBC count, RBC count, platelet count, creatinine, LDL cholesterol, and NfL, no relevant multicollinearity was present (all VIFs ≤ 1.35).

In this full model, platelet count (standardized β = 0.309, *p* = 0.005) and creatinine (β = 0.277, *p* = 0.016) were independently associated with higher serum sPDGFRβ levels. LDL cholesterol showed a positive but borderline association with sPDGFRβ (β = 0.189, B = 15.22, 95% CI 1.81 to 32.25, *p* = 0.079), whereas WBC count (*p* = 0.842), RBC count (*p* = 0.298), and NfL (*p* = 0.138) were not independently associated with sPDGFRβ (Table 2).

During backward elimination, WBC count was removed first, followed by RBC count and NfL. The final parsimonious model retained platelet count, creatinine, and LDL cholesterol. All three variables were independently and positively associated with serum sPDGFRβ: platelet count (β = 0.275, B = 0.132, 95% CI 0.036 to 0.227, *p* = 0.007), creatinine (β = 0.327, B = 0.290, 95% CI 0.112 to 0.468, *p* = 0.002), and LDL cholesterol (β = 0.238, B = 19.10, 95% CI 3.05 to 35.15, *p* = 0.020) (Table 2). Multicollinearity remained negligible in the final model (VIFs 1.027–1.046).

Neither cohort showed a significant linear correlation between sPDGFRβ and NIHSS; similarly, sPDGFRβ was not linearly associated with 3-month mRS in the Taizhou cohort (Tables 1 and 2). Together with the between-group differences described above (Fig. 1, A and C), these findings suggest that any relationship between circulating sPDGFRβ and global clinical severity may be weak or nonlinear and will require confirmation in larger cohorts.

### High-fat feeding suppressed interferon signaling and shifted oxidative-stress programs in cerebral pericytes

The replicated clinical association between LDL cholesterol and serum sPDGFRβ prompted us to examine cerebral pericyte responses to atherogenic lipid stress *in vivo*. Twenty-one *Apoe*-knockout mice were fed HFD or standard chow for 2 months. Serum lipid measurements from 21 *Apoe*-knockout mice confirmed the establishment of hypercholesterolemia. High-fat feeding increased both total and LDL cholesterol, but the LDL fraction responded disproportionately. Total cholesterol rose 1.27-fold (95% CI 1.15–1.41, *p* = 9.5 × 10⁻⁵) whereas LDL cholesterol rose 2.05-fold (95% CI 1.77–2.38, *p* = 6.2 × 10⁻⁹); in a mixed-effects model of log₂-transformed concentrations with a random intercept per animal, the analyte × diet interaction corresponded to a 1.61-fold steeper relative response of LDL cholesterol (95% CI 1.41–1.84, *p* = 4.7 × 10⁻⁷, adjusted for sex; exact permutation *p* = 8.6 × 10⁻⁶). Non-LDL cholesterol was unchanged (1.09-fold, *p* = 0.13), indicating that the hypercholesterolemia induced by high-fat feeding in this model is driven predominantly by the LDL compartment (Supplementary Fig. S1). Additionally, atherosclerotic plaques were detected in the aortic arch of HFD-fed but not control mice, confirming a diet-associated atherogenic phenotype (Supplementary Fig. S2).

A subset of eight mice (n=2 per sex × diet group) subsequently underwent single-cell RNA sequencing. Single-cell RNA sequencing yielded 104,032 cells after quality filtering and doublet removal, with major neurovascular and immune cell populations represented (Fig. 2, A and B). Pericytes were defined by high expression of canonical markers, including *Pdgfrb*, *Rgs5*, *Kcnj8*, and *Atp13a5*, together with low expression of endothelial (*Cldn5*, *Pecam1*), vascular smooth muscle/contractile (*Myh11*), and microglial (*Cx3cr1*, *C1qa*) markers (Fig. 2, B).

**Figure 2.**
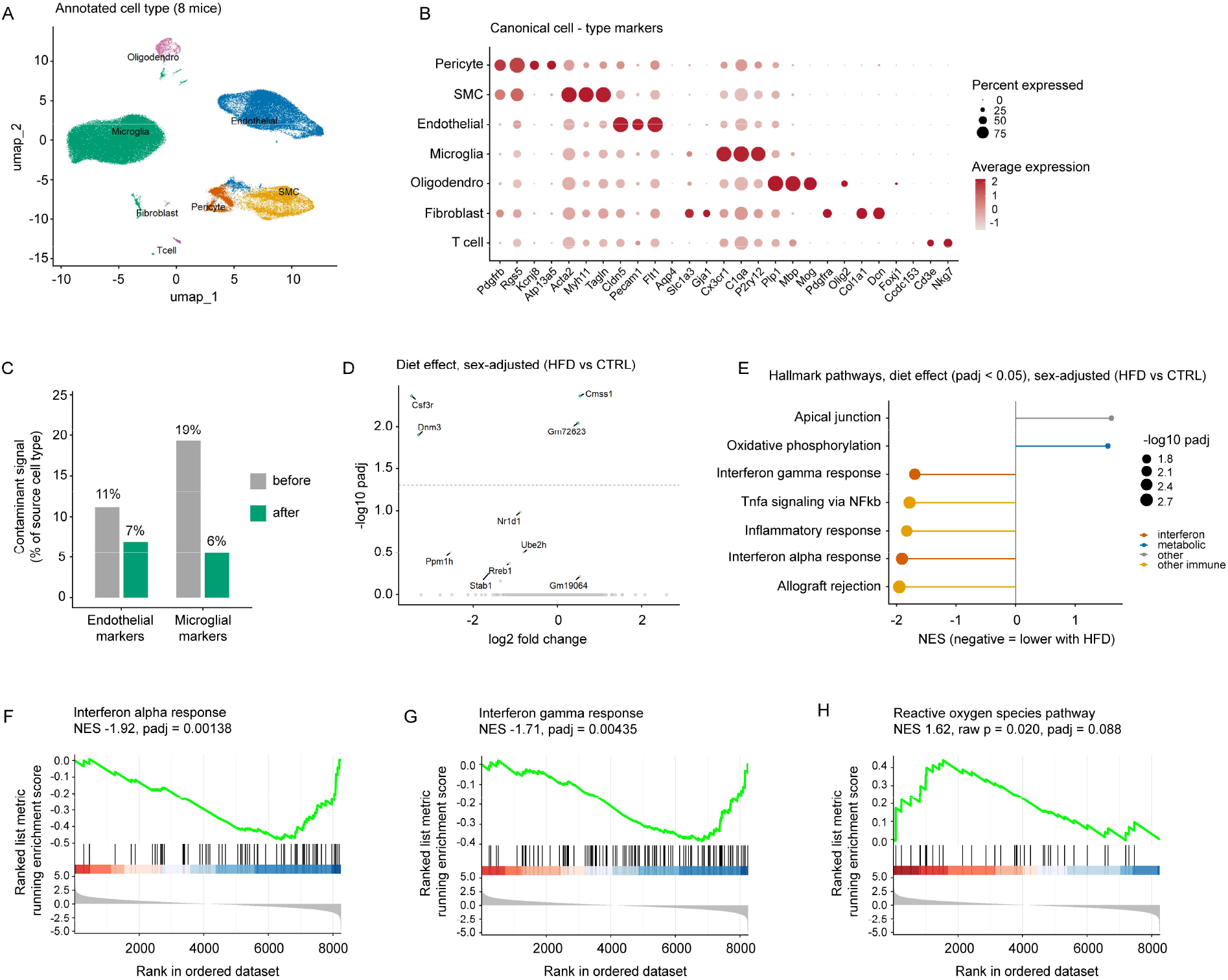
High-fat diet suppressed interferon-related programs and shifted oxidative-stress signaling in cerebral pericytes. Single-cell RNA sequencing was performed in 8 Apoe-knockout mice receiving control chow (CTRL) or high-fat diet (HFD; *n* = 2 per sex × diet group). UMAP shows major neurovascular and immune cell populations (A), and canonical marker expression was used for cell-type annotation (B). Pericyte purification and DecontX correction reduced residual endothelial and microglial marker signals, yielding 2588 pericytes for analysis (C). The volcano plot shows the sex-adjusted HFD effect in per-animal pseudobulk profiles (DESeq2, ∼ sex + diet) (D). Hallmark pathways with adjusted *p* < 0.05 are shown by normalized enrichment score (NES), with negative values indicating lower pathway activity with HFD (E). GSEA plots show IFN-α response (F), IFN-γ response (G), and reactive oxygen species response (H). The animal, rather than the individual cell, was the unit of biological replication.

Because interferon-stimulated genes are abundant in microglia and endothelial cells, we specifically addressed contamination before differential expression analysis. Of 4048 cells initially assigned to pericyte clusters, subclustering and per-cell filtering retained 2600 cells with pericyte-dominant signatures. DecontX further reduced the relative microglial marker burden from 19% to 6% and the endothelial marker burden from 11% to 7% (Fig. 2, C). After exclusion of 12 cells with estimated contamination > 50%, the final dataset comprised 2588 pericytes, with 172 to 491 cells contributed by each animal.

Per-animal pseudobulk analysis showed modest effects of HFD at the individual-gene level, with only 4 genes reaching genome-wide significance (adjusted *p* < 0.05; Fig. 2, D). *Cmss1* and *Gm72623* had relatively high baseline expression, whereas *Csf3r* and *Dnm3* had low baseline expression (baseMean < 25) and should be interpreted cautiously. *Nr1d1*, a circadian and metabolic regulator, was among the leading diet-responsive genes (log2FC = −0.88, adjusted *p* = 0.10).

At the pathway level, however, the sex-adjusted diet contrast revealed coordinated transcriptional changes. Seven Hallmark pathways reached adjusted *p* < 0.05 (Fig. 2, E). IFN-α response (Fig. 2, F; NES = −1.92, adjusted *p* = 1.4 × 10⁻³) and IFN-γ response (Fig. 2, G; NES = −1.71, adjusted *p* = 4.4 × 10⁻³) were the most strongly depleted pathways, and TNFα signaling via NF-κB and inflammatory response were also depleted. Oxidative phosphorylation and apical junction pathways were enriched (Fig. 2, E; all adjusted *p* < 0.05). Given the marked oxidative-stress response to LDL in cultured pericytes (Fig. 4), we specifically examined the Hallmark reactive oxygen species pathway *in vivo*; it showed a concordant positive shift (NES = 1.62, raw *p* = 0.020), although it did not survive correction for multiple testing (adjusted *p* = 0.088) (Fig. 2, H). All Hallmark pathways tested are listed in Supplementary table 5.

Reactome analysis showed a concordant pattern, with IFN-α/β signaling as the highest-ranked interferon-related pathway (NES = −1.93). These findings indicate that chronic HFD exposure is associated with coordinated suppression of interferon-related programs in cerebral pericytes, accompanied by a suggestive shift toward oxidative-stress signaling.

### The cerebral pericyte response to high-fat diet was sex-dependent

We next tested whether the pericyte response to HFD differed by sex using a sex-by-diet interaction model. Eight Hallmark pathways showed significant sex-dependent responses (Fig. 3, A; adjusted *p* < 0.05), including an immune/inflammatory arm comprising TNFα signaling via NF-κB (NES = −1.79, adjusted *p* = 2.3 × 10⁻³), IFN-γ response (NES = −1.81, adjusted *p* = 3.6 × 10⁻³), inflammatory response (NES = −1.62, adjusted *p* = 3.4 × 10⁻²), and IFN-α response (NES = −1.63, adjusted *p* = 3.9 × 10⁻²); a metabolic arm comprising cholesterol homeostasis (NES = −1.89, adjusted *p* = 5.1 × 10⁻³), oxidative phosphorylation (NES = +1.47, adjusted *p* = 3.9 × 10⁻²), and heme metabolism (NES = +1.53, adjusted *p* = 4.4 × 10⁻²); and a depleted androgen-response pathway (NES = −1.64, adjusted *p* = 3.6 × 10⁻²). A curated type I interferon (IFN-I) module showed a particularly strong interaction (Fig. 3, B; NES = −2.05, *p* = 3.9 × 10⁻⁴), with the leading-edge genes including *Ifit1*, *Ifitm3*, *Bst2*, *Irf7*, *Ifit3*, *Ifitm1*, and *Cxcl10*. For pathways with negative enrichment scores, the high-fat-diet-associated decrease was more pronounced in females than in males.

**Figure 3.**
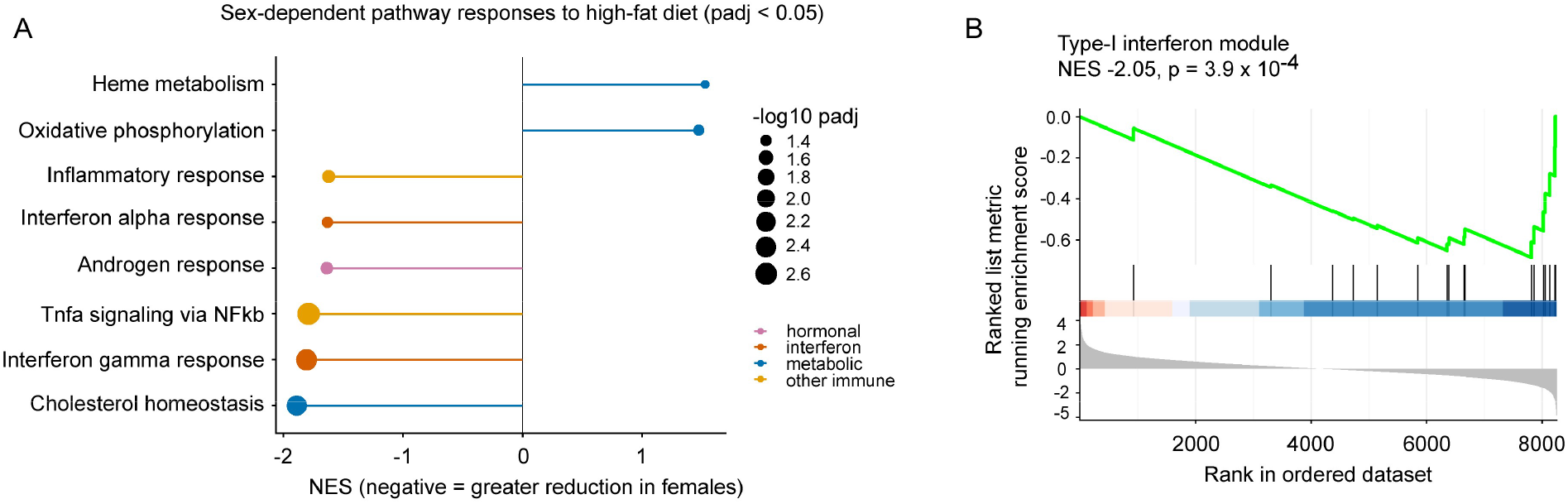
High-fat diet elicited sex-dependent transcriptional responses in cerebral pericytes. Per-animal pseudobulk profiles of 2588 decontaminated pericytes were modeled with a sex-by-diet interaction design (DESeq2, ∼ sex × diet; n = 2 animals per sex-by-diet group). Hallmark pathways with significant sex-dependent responses are ordered by normalized enrichment score (NES); negative NES values indicate a greater HFD-associated reduction in females than in males (A). GSEA of a curated IFN-I module against the sex-by-diet interaction ranking is shown in (B); leading-edge genes included *Ifit1*, *Ifitm3*, *Bst2*, *Irf7*, *Ifit3*, *Ifitm1*, and *Cxcl10*.

To determine whether residual microglial RNA could account for the IFN-I signal in pericytes, we repeated the enrichment analysis in the 50% of pericytes with the lowest estimated contamination. The IFN-I interaction was strengthened rather than attenuated (NES = −2.17, *p* = 1.3 × 10⁻⁵; Supplementary Fig. S3), arguing against residual microglial contamination as the primary explanation and supporting a sex-dependent pericyte interferon response to HFD.

### LDL cholesterol induced oxidative-stress and ferroptosis-associated programs and suppressed IFN-I signaling in human pericytes

To test whether LDL directly alters human pericytes, we exposed cultured pericytes to 2 mg/mL purified human LDL for 24 hours. mRNA sequencing identified 498 downregulated and 350 upregulated genes at |log2 fold change| ≥1 and FDR <0.01 (Fig. 4, A); randomly selected differentially expressed genes (DEGs) were validated by real-time PCR (Supplementary Fig. S4). *IFI35*, *IFIT1*, *SAMHD1* and *STAT1* were among the downregulated genes enriched in Gene Ontology terms related to cellular response to IFN-I and IFN-I signaling (Fig. 4, B; FDR < 0.01), paralleling the interferon suppression observed *in vivo* (Fig. 2, F and G). LDL also markedly reduced *LDLR* and *PCSK9* expression and strongly induced *PLIN2*, consistent with remodeling of cholesterol-homeostatic and lipid-storage pathways. Gene Ontology analysis showed induction of oxidative-stress (*GCLC*, *ABCC2*, *NGFR*, *PTGS2*, *PTGS1*, *DDIT3*, *NQO1*, *RBM11*, *GPX3* and *SRXN1*) and oxidoreductase programs (*HSD17B6*, *AIFM2*, *CYB5R4*, *AKR1B1*, *HSD17B2*, *NOX4*, *NQO2*, *AOC3*, *DMGDH*, *AKR1C2*, *NQO1*, *AKR1C1*, *AKR1C3*, *AKR1B10*, and *SRXN1*), together with changes in cytokine-related genes (*IL1β*, *IL6*, *IL11*, *TNFSF9*, *TNFSF11* and *NLRP10*) (Fig. 4, C; FDR < 0.01). Kyoto Encyclopedia of Genes and Genomes analysis further identified a ferroptosis-associated transcriptional response involving: *GCLC*, *GCLM*, *FTH1*, *FTL*, *HMOX1*, *SLC3A2*, and *SLC7A11* (Fig. 4, D; FDR < 0.01) ^31^. These data indicate oxidative and lipid stress but do not by themselves establish ferroptotic cell death.

**Figure 4.**
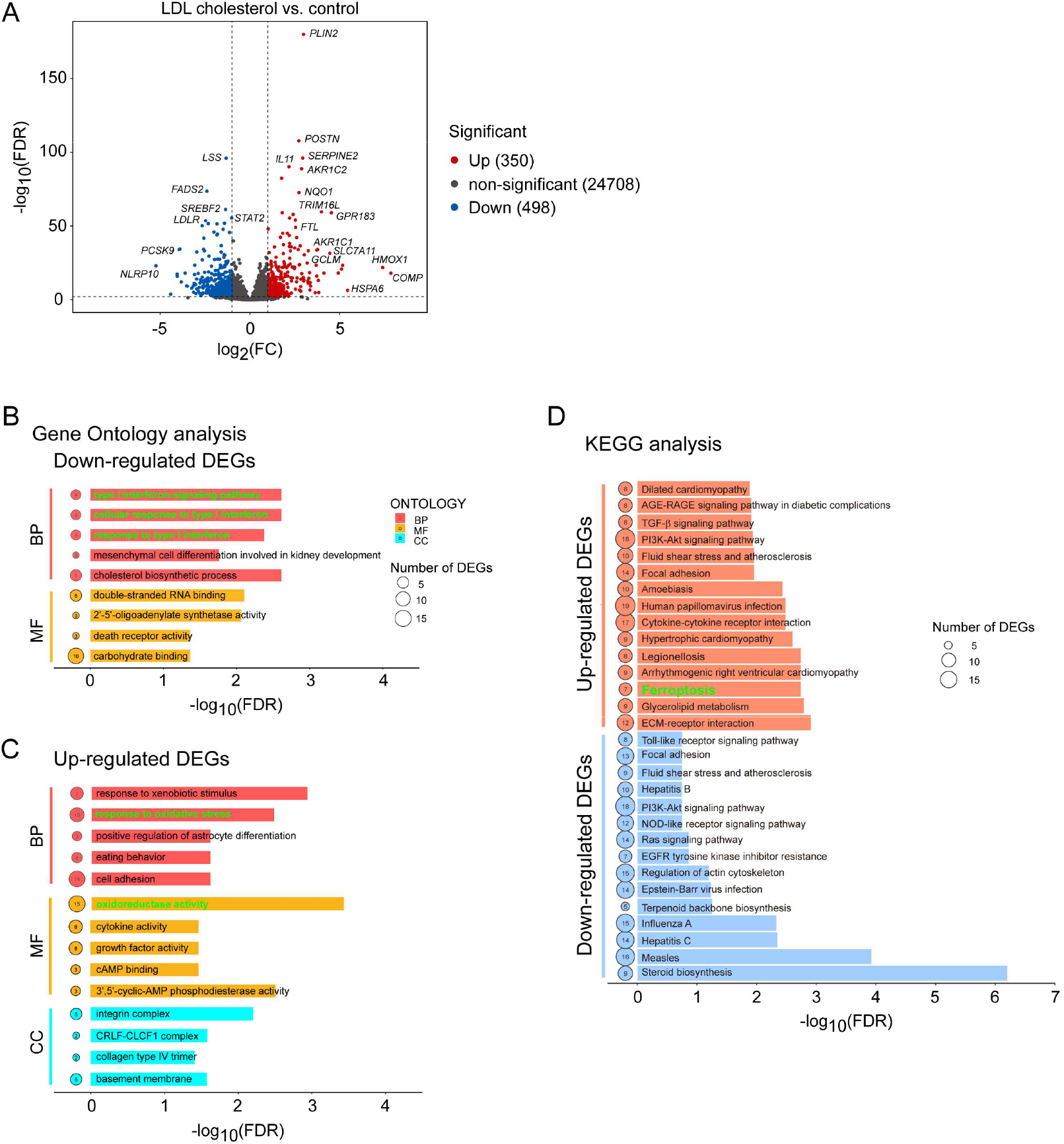
LDL cholesterol induced oxidative-stress and ferroptosis-associated transcriptional programs and suppressed IFN-I signaling in cultured human pericytes. Cultured human pericytes were exposed to 0 or 2 mg/mL purified human LDL for 24 hours, followed by mRNA sequencing. Differentially expressed genes were defined as |log2 fold change| ≥1 and FDR < 0.01 and are shown in the volcano plot (A). Downregulated genes were analyzed by Gene Ontology enrichment (B), and upregulated genes were analyzed for Gene Ontology terms related to oxidative stress, oxidoreductase activity, and cytokine activity (C). Kyoto Encyclopedia of Genes and Genomes analysis identified a ferroptosis-associated gene set among LDL-induced genes (D). n = 6 independent samples per group.

### *In vitro* LDL signatures mapped to the sex-dependent high-fat-diet response *in vivo*

We next tested whether LDL-induced transcriptional signatures in cultured human pericytes were represented in the mouse HFD response using ortholog-mapped GSEA. The LDL-repressed signature was significantly enriched at the negative interaction pole, indicating that these genes were reduced by HFD to a greater extent in females than in males (Fig. 5, A and C; NES = −1.82, *p* = 3.4 × 10⁻⁵). Leading-edge genes included an interferon core (*Stat1*, *Stat2*, *Irf7*, *Bst2*, *Ifitm1*, and *Ifitm3*) together with genes involved in cholesterol-biosynthetic feedback (*Srebf1*, *Fasn*, and *Ldlr*). Conversely, the LDL cholesterol - induced lipid/oxidative-stress signature was enriched at the positive interaction pole, indicating a greater HFD-associated increase in females (Fig. 5, A and D; NES = +1.66, *p* = 3.4 × 10⁻⁴). Neither signature was enriched in the sex-adjusted average diet effect (Fig. 5, A), indicating that cross-system correspondence was primarily related to the sex-dependent HFD response.

**Figure 5.**
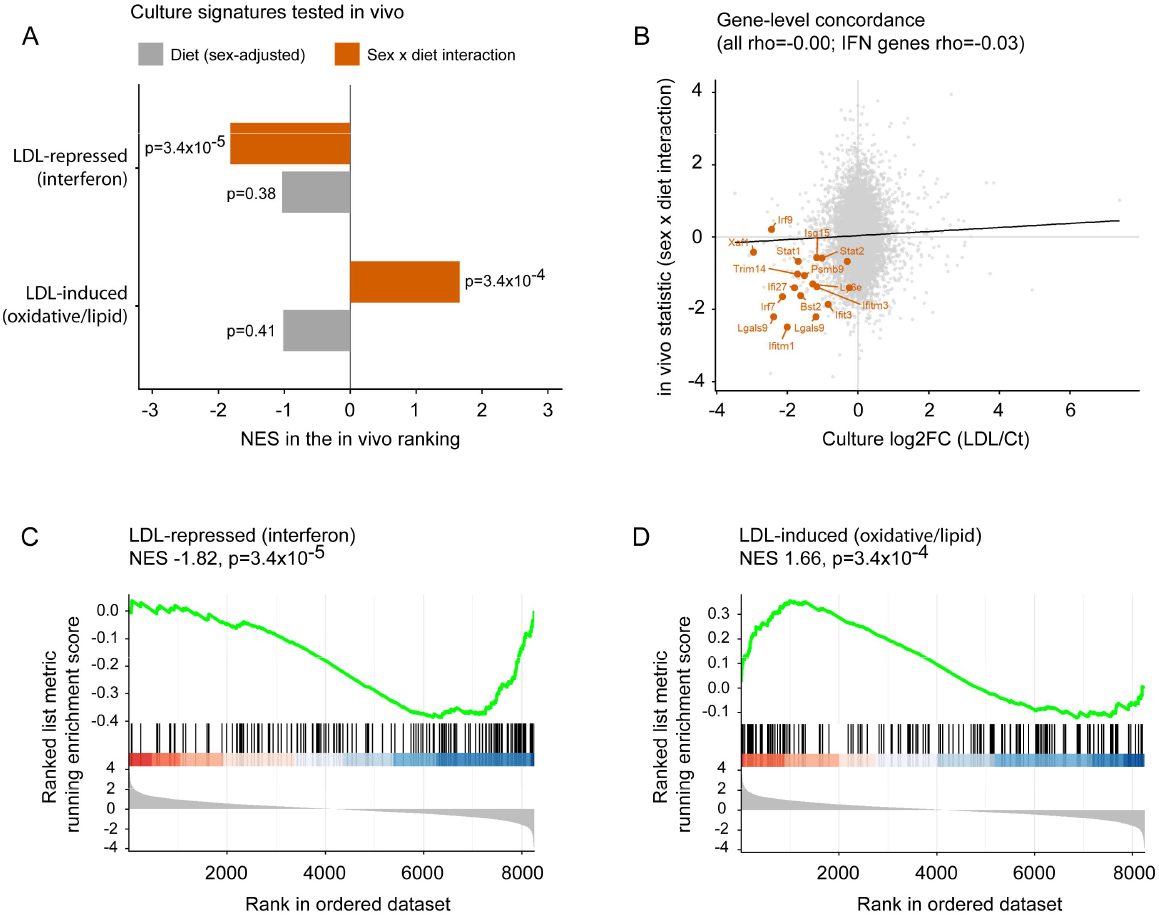
Cultured-pericyte LDL signatures aligned with the sex-dependent high-fat-diet response *in vivo*. LDL-induced and LDL-repressed signatures from cultured human pericytes (FDR < 0.05 and |log2 fold change| ≥ 1; n = 6 per group) were mapped to mouse orthologs and tested by GSEA against the sex-adjusted diet effect and sex-by-diet interaction. Neither signature was enriched in the average diet response, whereas both were strongly enriched in the sex-by-diet interaction (A), Gene-level concordance between the *in vitro* LDL response and *in vivo* interaction statistic is shown for shared orthologs (grey), with interferon-stimulated genes highlighted (orange) (B; Spearman correlation test). The LDL-repressed, interferon-dominated signature was enriched toward genes reduced by HFD to a greater extent in females than in males (C), whereas the LDL-induced lipid/oxidative-stress signature was enriched toward genes increased to a greater extent in females (D).

At the individual-gene level, LDL-induced fold changes in cultured pericytes showed essentially no global correlation with the *in vivo* sex-by-diet interaction statistic (Fig. 5B; Spearman ρ = 0.00), and the relationship remained weak when restricted to interferon-related genes (ρ = −0.03). Thus, concordance between the two systems emerged at the level of coordinated transcriptional programs rather than uniform gene-by-gene responses. Together, these analyses identify a reciprocal lipid-interferon axis in which LDL-associated lipid/oxidative programs and interferon suppression show sex-dependent relationships *in vivo*.

### LDL cholesterol reduced cell-surface PDGFRβ and increased sPDGFRβ release in human pericytes

We next examined the effects of LDL cholesterol on PDGFRβ at the protein level and assessed whether these changes were accompanied by cellular injury. Flow cytometry showed that LDL exposure reduced cell-surface PDGFRβ (Fig. 6, A - D). In the absence of NAC, the proportion of PDGFRβ-positive cells showed a significant LDL concentration effect (Fig. 6, C; Greenhouse–Geisser-corrected *p* = 0.036), with the lowest values observed at 2 mg/mL LDL. PDGFRβ fluorescence intensity showed a concordant decrease at 2 mg/mL LDL, although the overall concentration effect did not reach statistical significance (Fig. 6, D; *p* = 0.151).

**Figure 6.**
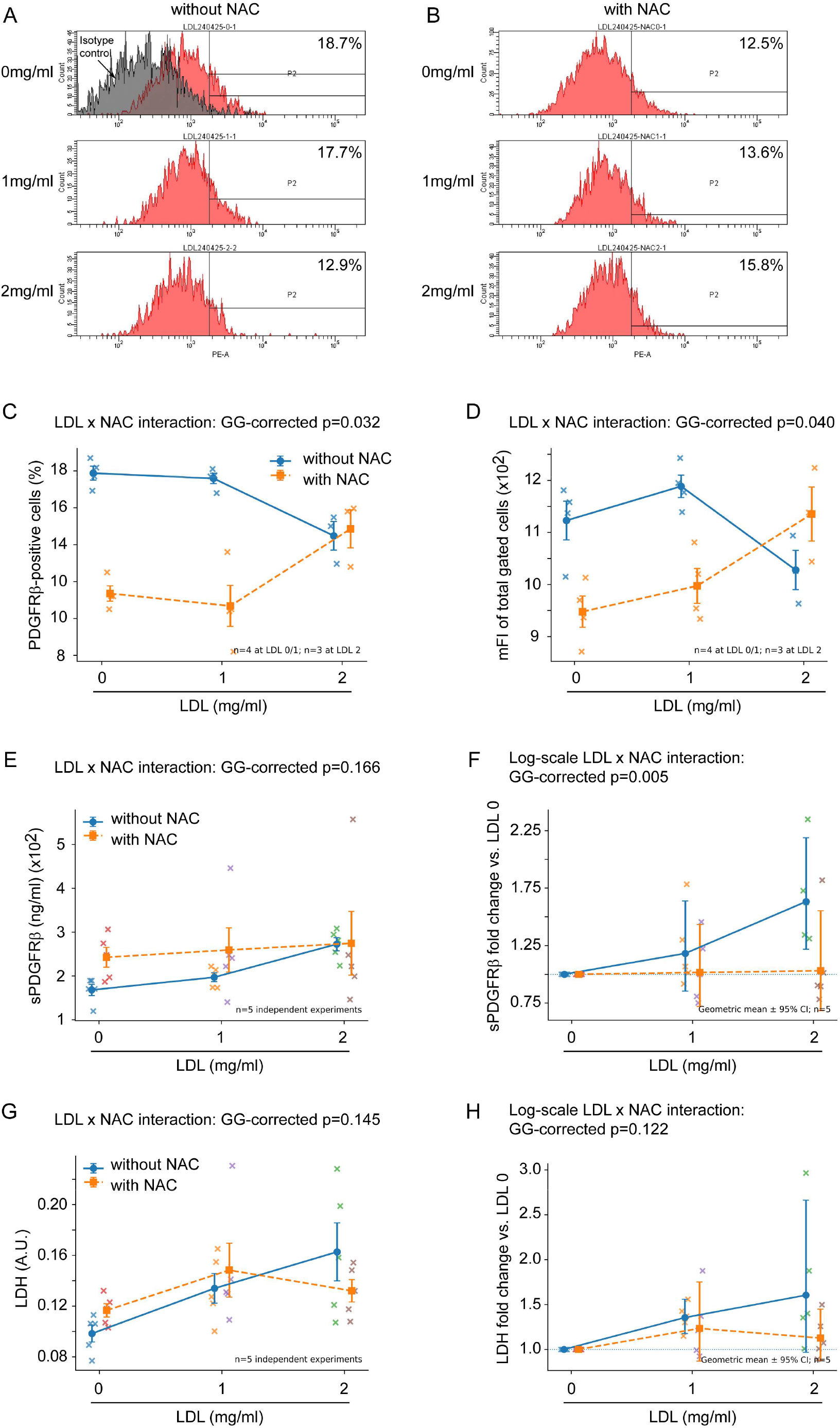
LDL cholesterol reduces cell-surface PDGFRβ and increases sPDGFRβ release in human brain pericytes, with modulation by N-acetyl-L-cysteine. (Figure legend followed on the next page) Flow cytometry was used to analyze cell-surface PDGFRβ in human brain pericytes exposed for 24 hours to 0, 1, or 2 mg/mL LDL cholesterol in the absence (A) or presence (B) of 2 mmol/L N-acetyl-L-cysteine (NAC). In the absence of NAC, LDL concentration significantly affected the proportion of PDGFRβ-positive cells (C; Greenhouse-Geisser-corrected *p*=0.036), with the lowest values at 2 mg/mL LDL, whereas the corresponding effect on mean fluorescence intensity (mFI) did not reach statistical significance (D; *p*=0.151). Four independent flow-cytometry experiments were performed at 0 and 1 mg/mL LDL; the 2 mg/mL condition was unavailable in one experiment, leaving three complete experiments for factorial analysis. Two-way repeated-measures ANOVA demonstrated significant LDL×NAC interactions for both the proportion of PDGFRβ-positive cells (C; *p*=0.032; partial η²=0.87) and PDGFRβ mFI (D; *p*=0.040; partial η²=0.82). Extracellular sPDGFRβ was measured in conditioned medium from five independent experiments. In the absence of NAC, LDL significantly increased sPDGFRβ concentrations (E; Greenhouse-Geisser-corrected *p*=0.005; partial η²=0.83), with sPDGFRβ at 2 mg/mL LDL significantly higher than untreated control after Holm correction (*p*=0.013). No significant LDL concentration effect was detected in NAC-treated cells (E; *p*=0.640), and the LDL×NAC interaction on the original concentration scale was not significant (*p*=0.166). Proportional sPDGFRβ responses were therefore additionally evaluated after normalization to the corresponding LDL 0 condition and logarithmic transformation, which demonstrated a significant LDL×NAC interaction (F; Greenhouse-Geisser-corrected *p*=0.005; partial η²=0.76). LDH activity was measured in conditioned medium from five independent experiments as an index of cytotoxicity. In the absence of NAC, LDH activity increased numerically with LDL concentration, but the overall LDL effect did not reach statistical significance (G; *p*=0.096); the LDL×NAC interaction was also not significant (*p*=0.145). LDH responses normalized to the corresponding LDL 0 condition likewise showed no significant interaction after logarithmic transformation (H; *p*=0.122). Individual symbols represent independent experiments. Data in C–E and G are shown as mean ± SEM; normalized data in F and H are shown as geometric mean with 95% confidence intervals. LDL concentration and NAC treatment were analyzed as within-experiment factors using repeated-measures ANOVA with Greenhouse-Geisser correction.

Consistent with reduced cell-surface PDGFRβ, LDL markedly increased extracellular sPDGFRβ release in five independent experiments. In the absence of NAC, mean sPDGFRβ increased from 168.0 ± 12.8 at 0 mg/mL LDL to 196.9 ± 10.1 at 1 mg/mL and 272.4 ± 15.1 at 2 mg/mL LDL (Fig. 6, E; repeated-measures ANOVA, Greenhouse–Geisser-corrected *p* = 0.0046; partial η² = 0.83). sPDGFRβ at 2 mg/mL LDL was significantly higher than untreated control after correction for multiple comparisons (Holm-adjusted *p* = 0.013). LDL exposure was also accompanied by a numerical increase in LDH activity, from 0.098 ± 0.007 at baseline to 0.134 ± 0.012 and 0.163 ± 0.023 at 1 and 2 mg/mL LDL, respectively (Fig. 6, G). However, the overall LDL effect on LDH did not reach statistical significance after Greenhouse–Geisser correction (*p* = 0.096). Thus, the effects of LDL on PDGFRβ surface availability and sPDGFRβ release were more pronounced than the accompanying change in LDH activity.

We next examined whether the antioxidant NAC modified these LDL responses. Significant LDL × NAC interactions were observed for both the proportion of PDGFRβ-positive cells (Fig. 6, C; Greenhouse–Geisser-corrected *p* = 0.032; partial η² = 0.87) and PDGFRβ fluorescence intensity (Fig. 6, D; *p* = 0.040; partial η² = 0.82). Whereas increasing LDL was associated with loss of cell-surface PDGFRβ in the absence of NAC, the concentration-response profile was shifted in the opposite direction in NAC-treated cells, indicating that NAC significantly modified the effect of LDL on cell-surface PDGFRβ (Fig. 6, C and D).

NAC also modified the proportional sPDGFRβ response to LDL. No significant LDL concentration effect was detected in NAC-treated cells on the original concentration scale (Fig. 6, E; *p* = 0.640), and the overall LDL×NAC interaction for absolute sPDGFRβ concentrations did not reach statistical significance (*p* = 0.166). Because absolute sPDGFRβ concentrations showed substantial between-experiment variation, proportional responses were additionally evaluated after logarithmic transformation. This analysis demonstrated a significant LDL×NAC interaction (Fig. 6, F; Greenhouse–Geisser-corrected *p* = 0.005; partial η² = 0.76). The geometric mean LDL2/LDL0 response was approximately 1.63-fold without NAC compared with 1.03-fold with NAC, corresponding to a ratio of responses of 0.63 (95% CI, 0.50 - 0.80; *p* = 0.006). In contrast, NAC did not significantly modify the LDL-associated change in LDH activity: the LDL×NAC interaction was nonsignificant on both the original scale (Fig. 6, G; *p* = 0.145) and after logarithmic transformation (Fig. 6, H; *p* = 0.122). Together, these findings indicate that LDL-induced changes in PDGFRβ surface availability and sPDGFRβ release were sensitive to antioxidant treatment and occurred in the absence of a comparably robust increase in LDH release.

## Discussion

Pericytes are critical for maintaining the structural and functional integrity of the cerebral microcirculation ^6^. In the present study, higher circulating sPDGFRβ was associated with greater neurological deficit and with NfL in patients with AIS, and LDL cholesterol was independently associated with sPDGFRβ in 2 separate clinical cohorts. Experimental studies provided complementary evidence for a direct lipid-pericyte interaction. High-fat feeding in *Apoe*-knockout mice and LDL exposure in cultured human pericytes converged on coordinated suppression of interferon-related transcriptional programs and activation of lipid/oxidative-stress responses. LDL cholesterol also directly reduced cell-surface PDGFRβ, and increased sPDGFRβ release. These effects were attenuated by an antioxidant, N-acetyl-L-cysteine. Together, the data support an LDL-pericyte axis as a potential link between atherogenic lipid stress and cerebral microvascular dysfunction in AIS.

Although serum sPDGFRβ was higher in patients with more severe neurological deficits, its linear associations with NIHSS and 3-month mRS were modest. This is not unexpected because sPDGFRβ likely reflects a specific neurovascular process, whereas NIHSS is strongly influenced by infarct location and lesion burden and mRS integrates subsequent complications, treatment, and rehabilitation. Serum sPDGFRβ may also be influenced by extracerebral sources, BBB permeability, and systemic protein clearance. In addition, different ELISA platforms were used in the 2 cohorts, so absolute sPDGFRβ concentrations should not be compared across sites. Nevertheless, the association between sPDGFRβ and NfL supports coupling between microvascular and neuronal damage after AIS ^30^. Notably, the median NIHSS scores in our cohorts were 3 and 4, substantially lower than the median NIHSS of 13 reported in the prior NfL study ^30^, suggesting that these circulating signals may already be detectable in relatively mild AIS. Thus, serum sPDGFRβ may be more informative as a marker of a specific neurovascular pathobiology than as a stand-alone measure of global stroke severity or prognosis.

The most reproducible clinical observation was the independent association between serum sPDGFRβ and LDL cholesterol in both AIS cohorts. This finding extends the potential vascular consequences of LDL cholesterol beyond large-artery atherosclerosis ^32^ to the cerebral microcirculation. Given that pericytes share some phenotypic characteristics with vascular smooth muscle cells ^24, 33^, we hypothesized that they could be a direct target of LDL cholesterol. Indeed, our experiment showed that native, non-oxidized human LDL cholesterol directly reduced PDGFRβ surface expression on cultured pericytes and increased PDGFRβ shedding and LDH release, and these changes were attenuated by NAC, supporting oxidative stress as one mediator. The transcriptomic response was consistent with this interpretation: LDL cholesterol upregulated transcription of *PLIN2* and multiple oxidative-stress/oxidoreductase genes and activated a ferroptosis-associated gene set. LDL cholesterol also reduced *LDLR* and *PCSK9* expression, consistent with remodeling of cholesterol-homeostatic pathways during lipid loading. Prior work showing that native LDL cholesterol alters pericyte calcium handling^34^ supports a direct effect, while indirect effects mediated through LDL-induced endothelial dysfunction remain plausible because endothelial cells and pericytes are tightly coupled within the neurovascular unit ^6, 35^.

An additional finding was the convergence between the transcriptional responses of cerebral pericytes to chronic high-fat feeding *in vivo* and to acute LDL cholesterol exposure *in vitro*, which was characterized by coordinated suppression of interferon-related pathways, including IFN-α/β signaling, and a positive shift in reactive oxygen species-related transcriptional programs. Exploratory sex-by-diet analyses suggested that the reduction in IFN-I-related signaling and reciprocal lipid/oxidative-stress activation were more pronounced in females than in males. This observation is potentially relevant to broader cardiovascular microvascular biology. Women are five times more likely than men to present with ischemia or myocardial infarction with no obstructive coronary arteries (14.9% vs. 3.5%) ^36^. Coronary microvascular dysfunction is one recognized mechanism of ischemia without obstructive coronary disease, and hypercholesterolemia has been associated with reduced coronary flow reserve ^37, 38^. These cardiac observations do not establish an equivalent mechanism in the brain, but they provide a clinical context in which lipid-driven microvascular responses may be sex-sensitive.

However, the correspondence between the *in vivo* and *in vitro* models was evident primarily at the pathway level rather than at the level of individual genes. Gene-by-gene changes showed essentially no global correlation between LDL-treated cultured pericytes and the HFD sex×diet interaction in mice, indicating that the two experimental systems do not reproduce an identical transcriptional response. Consistent with this distinction, the oxidative response in cultured cells was dominated by inducible stress-response genes, including *HMOX1*, *NQO1*, *GCLM*, and *SLC7A11*, whereas the *in vivo* signal involved more constitutive antioxidant and mitochondrial components with leading-edge genes of *Gpx4*, *Sod1*, *Prdx2*, *Ftl1*, *Ndufa6*. Thus, the two models appear to share the same general biological direction—enhanced oxidative/lipid stress accompanied by reduced interferon activity—while differing in the magnitude and molecular implementation of that response.

Although HFD-induced hypercholesterolemia in our *Apoe*-knockout mice was driven predominantly by the measured LDL cholesterol fraction, the lipoprotein profile of *Apoe*-deficient mice differs substantially from that of humans. In *Apoe*-knockout mice, cholesterol is carried not only in LDL particles but also in cholesterol-rich VLDL and remnant lipoproteins^39^. Thus, the *in vivo* model reflects chronic exposure to a broader atherogenic lipoprotein milieu. By contrast, the cell-culture experiments, in which human pericytes were exposed directly to purified LDL cholesterol, provide more specific evidence for an LDL-mediated pericyte response. Oxidized LDL, a highly atherogenic modified form of LDL ^32^, was not quantified in either experimental system. We therefore cannot determine to what extent LDL oxidation contributed to the observed effects. Nevertheless, the selective rise in LDL cholesterol, the development of aortic atherosclerotic plaques in HFD-fed *Apoe*-knockout mice, and the convergent oxidative-stress and interferon-related transcriptional responses observed across the *in vivo* and *in vitro* models together support a shared pericyte response to atherogenic lipid stress.

The biological significance of LDL-associated suppression of IFN-I signaling remains unresolved. A recent preclinical report indicates that endothelial IFN-I signaling can contribute to vascular normalization after ischemic brain injury: endothelial *Ifnar1* deletion exacerbated poststroke BBB disruption, whereas IFN-β administration has been reported to reduce acute barrier injury and inflammatory-cell infiltration ^40–42^. If a similar protective program operates in pericytes, LDL cholesterol-mediated attenuation of IFN-I signaling could weaken microvascular adaptation to ischemia. However, IFN-I effects are highly cell-, dose-, and time-dependent, and the present study does not establish a causal role for pericyte IFN-I signaling. Pericyte-specific manipulation of IFNAR1 in hyperlipidemic stroke models will be required to determine whether this pathway mediates LDL-associated microvascular vulnerability.

The dissociation between PDGFRβ-related changes and LDH release upon the challenge of LDL cholesterol suggests that the increase in extracellular sPDGFRβ is unlikely to represent merely passive release secondary to generalized cell lysis. LDL produced a robust increase in sPDGFRβ together with reduced cell-surface PDGFRβ, which was attenuated by antioxidant treatment, whereas LDH activity showed only a variable, statistically nonsignificant increase and no significant LDL×NAC interaction. Collectively, these observations are compatible with oxidative stress-dependent regulation of PDGFRβ surface availability and/or ectodomain release occurring before or in parallel with overt cytotoxic injury. Nevertheless, because LDH activity showed a numerical increase with LDL exposure, the present experiments cannot exclude a contribution of sublethal membrane injury or cell death to sPDGFRβ release. This study has several limitations. First, the clinical cohorts were modest in size and observational; serum sPDGFRβ is not brain-specific, and different ELISA platforms were used across cohorts. These factors limited statistical power to assess relationships between serum sPDGFRβ and global clinical scores such as NIHSS and mRS. Second, the mouse experiment included only 2 animals per sex-by-diet group, no circulating lipoprotein fractions or oxidized LDL were quantified, and the mice were not subjected to cerebral ischemia; the sex-dependent findings therefore require validation in larger cohorts and experimental stroke models. Third, the *in vitro* experiments used a conditionally immortalized human brain pericyte line and an acute 2 mg/mL LDL exposure, which does not reproduce the complexity or duration of human dyslipidemia. Fourth, lipid peroxidation, ferroptotic cell death, and LDL uptake were not measured directly, and the *in vivo* study did not validate PDGFRβ shedding, capillary perfusion, or BBB function at the protein or physiological level. Finally, cross-species concordance emerged at the level of conserved transcriptional programs rather than individual genes.

In summary, two independent AIS cohorts linked circulating sPDGFRβ to LDL cholesterol, and complementary experimental models showed that atherogenic lipid exposure can reprogram cerebral pericytes and that LDL directly alters PDGFRβ homeostasis and oxidative-stress-associated transcriptional programs in human brain pericytes. Exploratory sex-dependent suppression of IFN-I and reciprocal activation of lipid/oxidative programs *in vivo* further suggest that sex may modify this response. These findings extend the vascular consequences of atherogenic lipids from large arteries to the cerebral microcirculation and motivate prospective studies of whether intensive lipid lowering preserves pericyte function and microvascular reperfusion after ischemic stroke.

## List of abbreviations

*AIS*: acute ischemic stroke
*BBB*: blood-brain barrier
*DEGs*: differentially expressed genes
*GO*: Gene Ontology
*KEGG*: Kyoto Encyclopedia of Genes and Genomes
*LDH*: lactate dehydrogenase
*LDL*: low-density lipoprotein
*mFI*: mean fluorescence intensity
*mRS*: modified Rankin scale
*NAC*: N-acetyl-l-cysteine
NfL: neurofilament light chain
*NIHSS*: National Institutes of Health stroke scale
*PDGFRβ*: platelet-derived growth factor receptor β
ROS: reactive oxygen species
sPDGFRβ: soluble platelet-derived growth factor receptor-β
VLDL: very-low-density lipoprotein

## CRediT authorship contribution statement

**Yang Liu:** Conceptualization, Funding acquisition, Investigation, Methodology, Data curation, Formal analysis, Supervision, Writing – original draft, Writing – review and editing. **Feng Wang:** Funding acquisition, Project administration, Investigation, Data curation, Formal analysis, Writing – original draft. **Wenqiang Quan:** Funding acquisition, Project administration, Investigation, Methodology, Data curation, Formal analysis, Writing – original draft. **Gang Wu:** Investigation, Methodology, Data curation, Formal analysis, Writing – original draft. **Jing Yang**: Investigation, Methodology, Data curation, Formal analysis. **Xu Wang:** Investigation, Methodology, Data curation, Formal analysis. **Qian Cao:** Investigation, Methodology, Data curation, Formal analysis. **Wenlin Hao:** Investigation, Methodology, Data curation, Formal analysis. **En Wang:** Investigation, Data curation, Formal analysis, Resources. **Tomomi Furihata:** Resources. **Weihai Xu:** Conceptualization, Writing – review and editing. **Dong Li:** Resources, Supervision. **Shaofa Ke:** Conceptualization, Funding acquisition, Resources, Supervision.

## Acknowledgements

Not applicable

## Ethical considerations

The clinical study protocol was approved by the Ethics Committees of Taizhou Hospital, Zhejiang Province (Registration number: K20181204) and Tongji Hospital, Shanghai, China (Registration number: 2021-LCYJ-011). The animal study was approved by the Ethics Committee for Animal Welfare at Taizhou Hospital, Zhejiang Province (Registration number: tzy-2024006).

## Consent to Participate

Written consent was provided by patients or their legal guardians.

## Consent for Publication

Not applicable.

## Declaration of generative AI and AI-assisted technologies

During preparation of this manuscript, Claude (Anthropic) and ChatGPT (OpenAI) were used to assist with language editing and structural refinement. After using these tools, the author(s) reviewed and edited the content and take full responsibility for the content of the published work.

