## Supplementary figures and tables for "Low-Density Lipoprotein Cholesterol and Cerebral Pericyte Dysfunction in Acute Ischemic Stroke: Clinical and Experimental Evidence"

### Supplemental Material

1. Supplementary Figure S1. High-fat diet raises LDL cholesterol disproportionately to total cholesterol in both sexes.
2. Supplementary Figure S2. High-fat diet induces atherosclerotic lipid deposition in Apoe−/− mice.
3. Supplementary Figure S3. Residual contamination does not account for the interferon findings in pericytes.
4. Supplementary Figure S4, Treatment with LDL cholesterol altered gene transcription in cultured pericytes involved in oxidative stress, ferroptosis and type I interferon signaling.

5. Supplementary Table 1, List of primer sequences used for quantitative PCR.
6. Supplementary Table 2, Characteristics of patients from Taizhou Hospital with different NIHSS scores at admission.
7. Supplementary Table 3, Multivariable logistic regression analysis of risk factors for NIHSS $\geq$ 4 at admission.
8. Supplementary Table 4, Characteristics of patients from Taizhou Hospital with different functional recovery at 3 months.
9. Supplementary Table 5, Hallmark pathways tested in cerebral pericytes of Apoe−/− mice fed with high-fat diet compared with standard chow (Excel table)

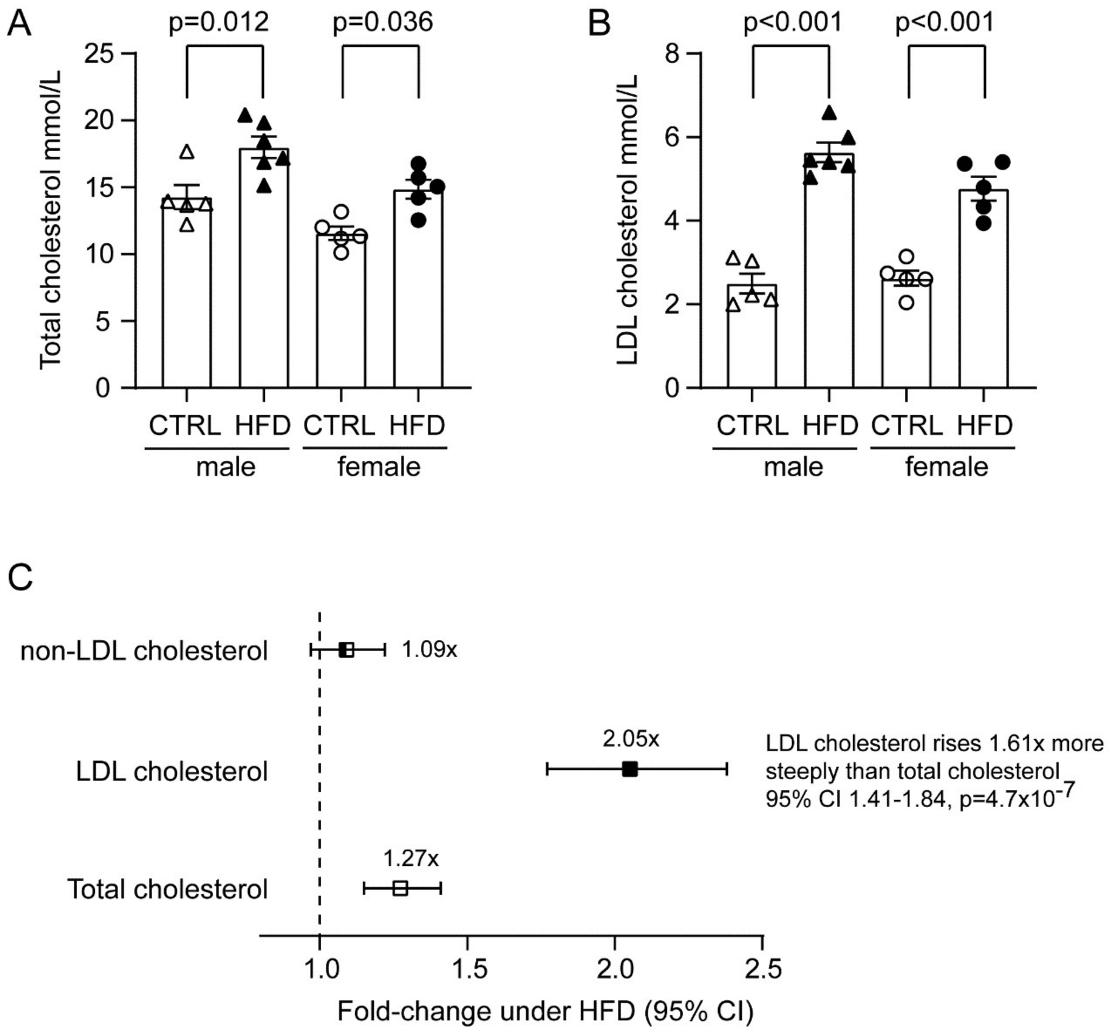


**Supplementary Figure S1. High-fat diet raises LDL cholesterol disproportionately to total cholesterol in both sexes.**

(A, B) Total cholesterol (A) and LDL cholesterol (B) in male (triangles) and female (circles) mice after high-fat diet (HFD, filled symbols) or standard chow (CTRL, open symbols). Bars show the group mean ± SEM with individual animals overlaid; n = 5 (male CTRL), 6 (male HFD), 5 (female CTRL) and 5 (female HFD). p values are from two-way ANOVA with diet and sex as factors followed by Tukey's multiple comparisons test.

(C) Fold-change of total, LDL and non-LDL cholesterol (total minus LDL) under HFD, pooled across sexes. Symbols show the point estimate and horizontal lines the 95% confidence interval, derived from linear models on $\log_2$-transformed concentrations adjusted for sex (n = 21 animals); the dashed line at 1.0 indicates no change. LDL cholesterol rose 2.05-fold (95% CI 1.77–2.38) and total cholesterol 1.27-fold (1.15–1.41), whereas non-LDL cholesterol was unchanged (1.09-fold, 0.97–1.22, p = 0.13). The LDL-C fold-change exceeded that of total cholesterol by a factor of 1.61 (95% CI 1.41–1.84, $p = 4.7 \times 10^{-7}$; analyte × diet interaction of a linear mixed-effects model on $\log_2$ concentrations with a random intercept per animal), indicating that hypercholesterolemia under high-fat feeding is carried predominantly by the LDL fraction.

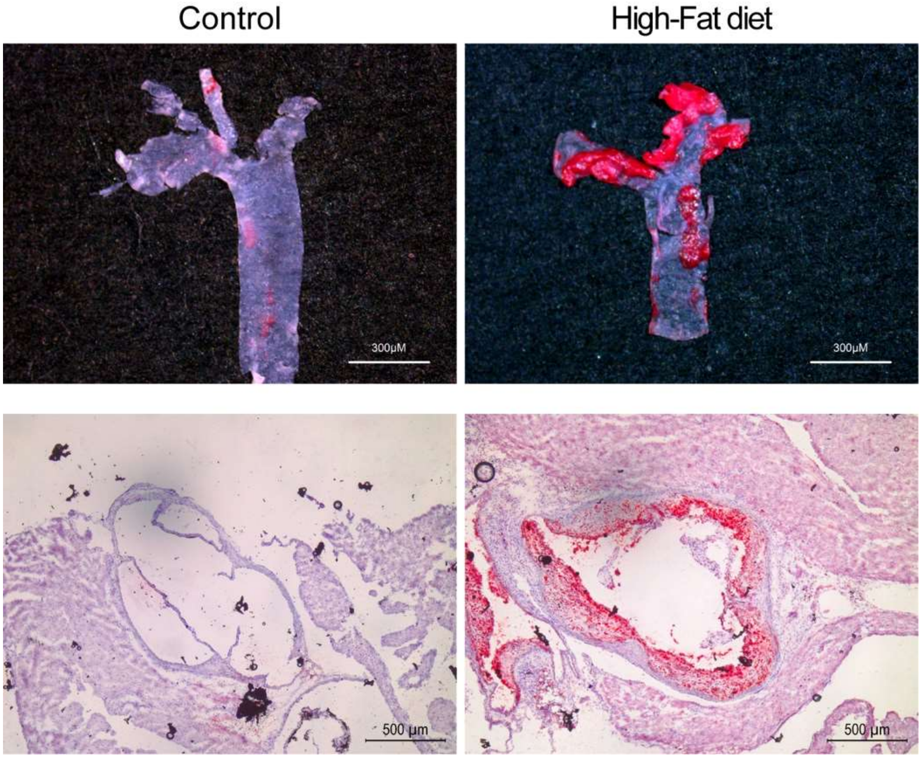


**Supplementary Figure S2. High-fat diet induces atherosclerotic lipid deposition in Apoe−/− mice.** Representative Oil Red O staining of the aortic arch from Apoe−/− mice fed a control diet (left panels) or a high-fat diet (right panels). Upper panels show en face preparations of the aortic arch, in which Oil Red O-positive lipid-rich atherosclerotic lesions are markedly increased following high-fat feeding. Lower panels show cross-sections of the aorta stained with Oil Red O and counterstained with hematoxylin, demonstrating prominent lipid accumulation and atherosclerotic plaque formation in high-fat diet-fed mice compared with controls. Red staining indicates neutral lipid deposition.

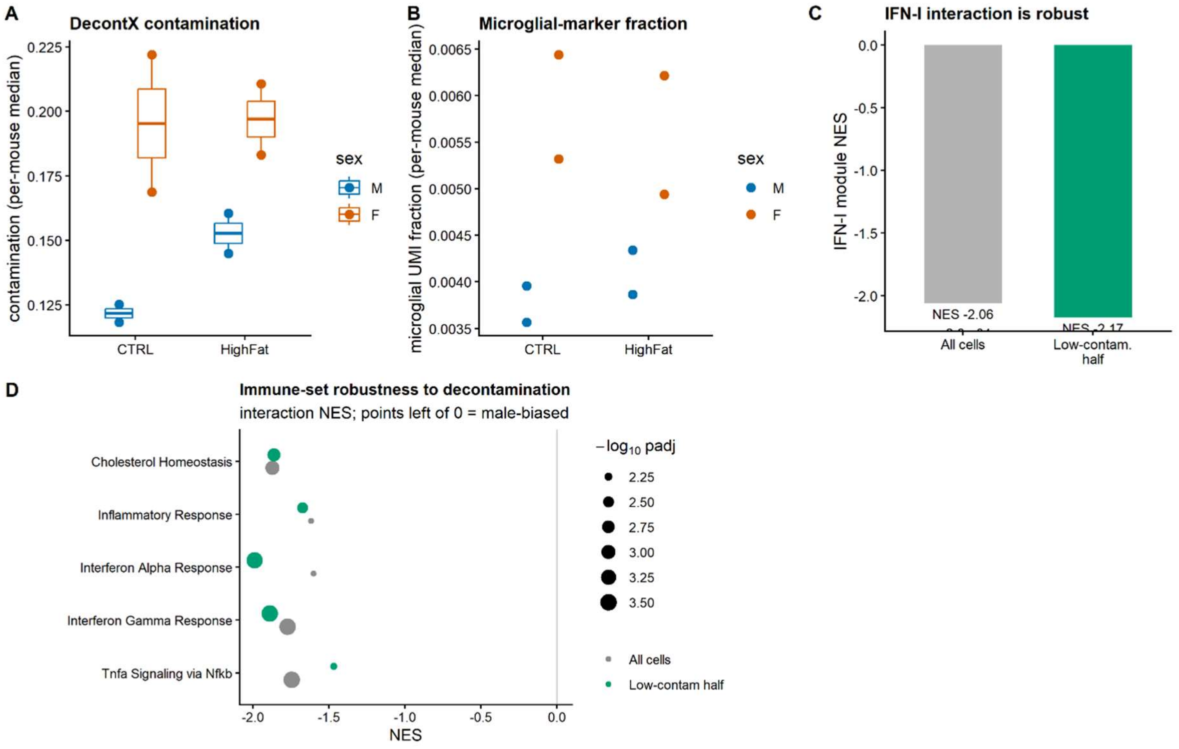


**Supplementary Figure S3. Residual contamination does not account for the interferon findings in pericytes.**

(A) Estimated ambient contamination (DecontX) in the final pericyte set, shown as the per-animal median for each sex × diet group (n = 2 animals per group). Contamination was comparable between diets (control 0.173 vs high-fat 0.186) and marginally higher in females than males (0.194 vs 0.139), i.e. opposite in direction to the male-biased interferon effect.

(B) Median fraction of UMIs assigned to microglial marker genes (*C1qa*, *C1qb*, *C1qc*, *Cx3cr1*, *Hexb*, *Ctss*, *P2ry12*, *Csf1r*, *Tmem119*) in the same cells, before decontamination, shown per animal by group. The microglial burden did not track the sex × diet effect (male high-fat, the group required to generate a spurious sex×diet interferon interaction, was among the lowest).

(C) Enrichment of the curated type-I interferon module against the sex × diet interaction, computed on all pericytes and, separately, on the half of cells with the lowest estimated contamination. Restricting the analysis to the least-contaminated cells strengthened rather than weakened the enrichment (NES −2.05 → −2.17; $p = 2.6 \times 10^{-4} \rightarrow 1.3 \times 10^{-5}$), as expected for a genuine signal and contrary to what contamination-driven enrichment would produce.

(D) The same test applied to the immune Hallmark gene sets enriched in the interaction (interferon-alpha response, interferon-gamma response, TNFα signaling via NFκB, inflammatory response), with cholesterol homeostasis included as a metabolic control that is not expected to be contamination-sensitive. Points show the normalized enrichment score for all pericytes (grey) and for the low-contamination half (green); point size indicates $-\log_{10}$ adjusted p. Enrichment was retained for interferon-alpha, and interferon-gamma, whereas TNFα signaling was weakened.

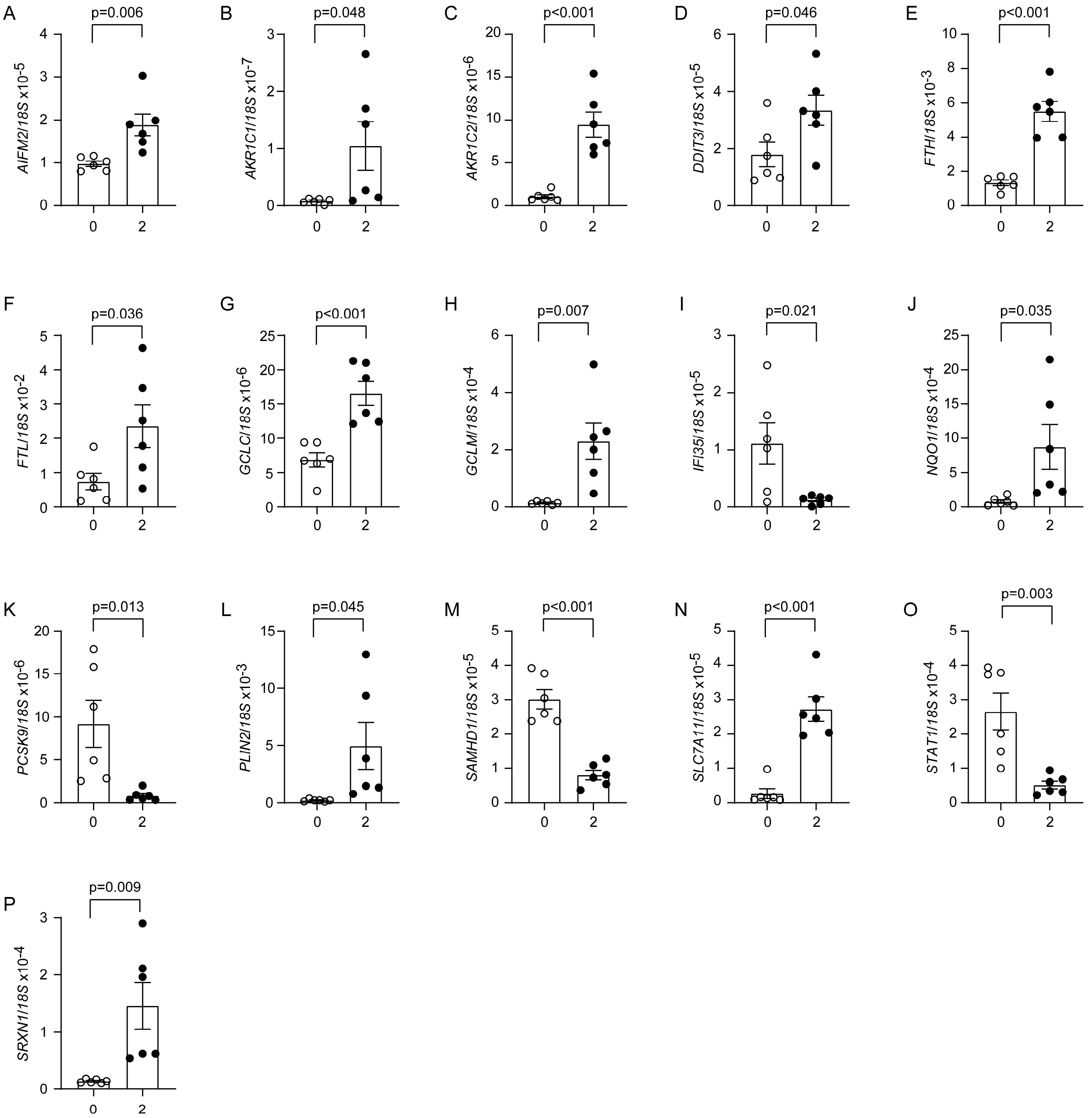


**Supplementary Figure S4, Treatment with LDL cholesterol altered gene transcription in cultured pericytes involved in oxidative stress, ferroptosis and type I interferon signaling.**

Cultured human brain pericytes (HBPC ci37) were treated with 0 and 2 mg/ml purified human LDL cholesterol for 24 hours. Total RNA was isolated and reverse transcribed. Different gene transcripts were measured with real-time PCR. *t*-test, $n = 6$ per group from 6 independent experiments.

Supplementary Table 1, List of primer sequences used for quantitative PCR

| Oligo Name | Sequence (5' → 3') |
|---|---|
| hPLIN2_for | CCGTTGCAGTTGATCCACAA |
| hPLIN2_rev | GTCTTCACACCGTTCTCTGC |
| hPCSK9_for | ACAGAGTGGGACATCACAGG |
| hPCSK9_rev | ACCAGGCCTCATTGATGACA |
| hAKR1C2_for | GGAGTTTCACCCTCTGGGAA |
| hAKR1C2_rev | GCTCCAGAAACTCTGACCCT |
| hNQO1_for | ATGGAAGAAACGCCTGGAGA |
| hNQO1_rev | GTCAGTTGGGATGGACTTGC |
| hFTL_for | AGAAGATGGGTGACCACCTG |
| hFTL_rev | CTATTGGCTGGAGGGAGAGG |
| hAKR1C1_for | GTTACCACTGCGTACCACAC |
| hAKR1C1_rev | GTTATGCGAGCGGTGAACTT |
| hGCLM_for | CCCAGATTTGGTCAGGGAGT |
| hGCLM_rev | CTGTGCAACTCCAAGGACTG |
| hDDIT3_for | TGAAAGGAAAGTGGCACAGC |
| hDDIT3_rev | CTGGGAAAGGTGGGTAGTGT |
| hIFI35_for | CTAGGGATGGAGTGGCTCAG |
| hIFI35_rev | CCCGTTCACATACGGAGAGA |
| hSAMHD1_for | CAGCAGATTTCCCAGCAGTC |
| hSAMHD1_rev | ACAGGATGGCAGACATTCCA |
| hSRXN1_for | GCTTGTCCAGTCCACTCTCT |
| hSRXN1_rev | GCTACTGCAAGTCTGGTGTG |
| hSTAT1_for | CTGGCACCAGAACGAATGAG |
| hSTAT1_rev | GAGGTCGTCTCGAGGTCAAT |
| hGCLC_for | AATGGGCAATTGCTGTCTCC |
| hGCLC_rev | GGGAGATGCAGCACTCAAAG |
| hAIFM2_for | CCGACAGTGCCTGATTTGAG |
| hAIFM2_rev | TCCTTCATGTCCACCAGCAT |
| hFTH1_for | TGAGGAGAGGGAACATGCTG |
| hFTH1_rev | TTGTCAGTGGCCAGTTTGTG |
| hSLC7A11_for | AACCGAAGGCCAGAGAATCA |
| hSLC7A11_rev | AGGTTCAGGACCTCGAATGG |

Supplementary Table 2, Characteristics of patients from Taizhou Hospital with different NIHSS scores at admission

| **Variables**[1] | Total (n = 97) | NIHSS score <4 (n =48) | NIHSS score ≥4 (n=49) | *P* value |
|---|---|---|---|---|
| **sPDGFRβ, ng/L** | 1664.61 ± 357.05 | 1577.86 ± 360.38 | 1745.75 ± 339.80 | 0.020 |
| **Demographic characteristics** | | | | |
| Female, n (%) | 42 (43.3) | 18 (37.5) | 24 (49.0) | 0.254 |
| Age, years | 67.90 ± 11.07 | 65.52 ± 10.58 | 69.94 ± 11.13 | 0.048 |
| **Risk factors for stroke** | | | | |
| Atrial fibrillation, n (%) | 8 (8.2) | 4 (8.3) | 4 (8.2) | 0.976 |
| Hypertension, n (%) | 49 (50.5) | 26 (54.2) | 23 (46.9) | 0.477 |
| Diabetes, n (%) | 18 (18.6) | 9 (18.8) | 9 (18.4) | 0.961 |
| Coronary heart disease, n (%) | 16 (16.5) | 8 (16.7) | 8 (16.3) | 0.964 |
| Previous stroke, n (%) | 4 (4.1) | 1 (2.1) | 3 (6.1) | 0.317 |
| Smoking, n (%) | 38 (39.2) | 20 (41.7) | 18 (36.7) | 0.619 |
| Drinking, n (%) | 15 (15.5) | 6 (12.5) | 9 (18.4) | 0.424 |
| **Laboratory findings** | | | | |
| White blood cells, $10^9$/L | 7.20 (5.50, 8.45) | 6.50 (5.23, 7.98) | 8.10 (5.80, 9.60) | 0.006 |
| Monocytes, $10^9$/L | 0.40 (0.30, 0.50) | 0.40 (0.30, 0.50) | 0.40 (0.30, 0.50) | 0.635 |
| Neutrophils, $10^9$/L | 4.60 (3.40, 6.00) | 4.10 (3.05, 5.18) | 5.40 (3.70, 7.20) | < 0.001 |
| Eosinophil, $10^9$/L | 0.09 (0.05, 0.20) | 0.10 (0.05, 0.20) | 0.08 (0.05, 0.24) | 0.274 |
| Basophil, $10^9$/L | 0.02 (0.01, 0.03) | 0.02 (0.01, 0.03) | 0.02 (0.02, 0.03) | 0.773 |
| Lymphocytes, $10^9$/L | 1.70 (1.25, 2.00) | 1.80 (1.33, 2.18) | 1.60 (1.20, 1.95) | 0.351 |
| Platelets, $10^9$/L | 227.09 ± 55.65 | 218.29 ± 51.97 | 230.98 ± 66.16 | 0.297 |
| Red blood cells, $10^9$/L | 4.42 ± 0.45 | 4.48 ± 0.40 | 4.30 ± 0.59 | 0.081 |
| Hemoglobin, g/L | 135.50 (126.75, 145.25) | 139.50 (130.25, 148.75) | 135.00 (125.50, 144.50) | 0.083 |
| C-reactive protein, mg/L | 2.40 (1.23, 6.20) | 2.45 (1.20, 5.40) | 2.00 (1.30, 6.25) | 0.442 |
| Fibrinogen, g/L | 3.09 (2.69, 3.59) | 3.16 (2.93, 3.57) | 3.12 (2.65, 4.01) | 0.707 |
| D-dimer, mg/L | 0.49 (0.28, 1.21) | 0.37 (0.28, 0.66) | 0.73 (0.34, 1.61) | 0.014 |
| Cr, µmol/L | 65.00 (53.00, 79.00) | 65.00 (53.75, 77.50) | 62.00 (53.00, 72.00) | 0.549 |
| BUN, mmol/L | 4.94 (4.14, 5.92) | 4.85 (4.47, 5.68) | 4.87 (4.85, 5.77) | 0.718 |
| UA, µmol/L | 289.00 (246.00, 351.50) | 302.00 (248.75, 359.25) | 275.00 (240.50, 314.50) | 0.248 |
| Glucose, mmol/L | 6.50 (5.58, 7.85) | 6.65 (5.60, 8.05) | 6.40 (5.36, 7.50) | 0.948 |
| HbA1c, % | 6.10 (5.88, 6.73) | 6.00 (5.75, 7.85) | 6.10 (5.90, 6.70) | 0.307 |
| Homocysteine, µmol/L | 11.75 (10.00, 14.75) | 12.15 (10.27, 16.18) | 10.80 (9.50, 12.40) | 0.279 |
| GPT, unit/L | 14.00 (11.00, 19.25) | 14.50 (10.00, 19.25) | 14.00 (12.00, 19.50) | 0.857 |
| GOT, unit/L | 20.00 (17.00, 25.00) | 18.50 (16.00, 23.25) | 20.00 (18.00, 24.00) | 0.151 |
| TBil, µmol/L | 13.55 (11.10, 17.85) | 11.90 (9.55, 15.90) | 14.70 (11.75, 20.55) | 0.027 |
| TG, mmol/L | 1.17 (0.86, 1.51) | 1.08 (0.92, 1.77) | 1.24 (0.85, 1.63) | 0.281 |
| TC, mmol/L | 4.48 ± 0.87 | 4.47 ± 0.85 | 4.49 ± 0.89 | 0.880 |
| HDL, mmol/L | 1.27 ± 0.25 | 1.26 ± 0.25 | 1.28 ± 0.25 | 0.743 |
| LDL, mmol/L | 2.53 ± 0.69 | 2.52 ± 0.67 | 2.55 ± 0.71 | 0.818 |
| **PDGF-BB, ng/L** | 495.63 (280.96, 698.06) | 475.00 (291.23, 569.86) | 496.36 (250.91, 737.44) | 0.690 |
| **Stroke onset – to – Blood collection time (h)** | 24.00 (7.25, 48.00) | 26.00 (13.50, 54.00) | 24.00 (7.00, 69.00) | 0.252 |
| **Thrombolysis therapy** | 9 (9.3) | 4 (8.3) | 5 (10.2) | 0.751 |

**Note:**

[1]Continuous variables with normal distribution were expressed as mean ± SD and non-normally distributed data were expressed as median (P25, P75). Categorical variables were presented as frequencies.

BUN, blood urea nitrogen; Cr, serum creatinine; GPT, glutamic pyruvic transaminase; GOT, glutamic oxaloacetic transaminase; HbA1c, Hemoglobin A1c; HDL, high density lipoprotein; LDL, low density lipoprotein; NIHSS, National Institute of Health stroke scale; TBil, Total bilirubin; TC, total cholesterol; TG, triglyceride; PDGF-BB, Platelet-derived growth factor-BB; UA, uric acid.

Supplementary Table 3, Multivariable logistic regression analysis of risk factors for NIHSS ≥ 4 at admission

| Independent variables | *P* value | OR | 95% CI |
|---|---|---|---|
| Age, years | 0.110 | 1.040 | - |
| White blood cells, $10^9$/L | 0.005 | 1.369 | 1.102 – 1.702 |
| Neutrophils, $10^9$/L | 0.881 | 0.993 | - |
| Red blood cells, $10^9$/L | 0.048 | 0.361 | 0.132 – 0.990 |
| Hemoglobulin, g/L | 0.845 | 0.999 | - |
| D-dimer, mg/L | 0.358 | 0.829 | - |
| Total bilirubin, μmol/L | 0.018 | 1.119 | 1.020 – 1.228 |
| **sPDGFRβ, ng/L** | 0.028 | 1.002 | 1.000 – 1.003 |

**Note:** OR, odds ratio; CI, confidence interval

Supplementary Table 4, Characteristics of patients from Taizhou Hospital with different functional recovery at 3 months

| Variables[1] | Total (n = 97) | mRS <2 (n =38) | mRS ≥2 (n=59) | *P* value |
|---|---|---|---|---|
| **sPDGFRβ, ng/L** | 1664.61 ± 357.05 | 1581.19 ± 350.47 | 1715.15 ± 356.48 | 0.072 |
| **Demographic characteristics** | | | | |
| Female, n (%) | 42 (43.3) | 15 (39.5) | 27 (45.8) | 0.542 |
| Age, years | 67.90 ± 11.07 | 67.82 ± 11.84 | 67.71 ± 10.58 | 0.964 |
| **Risk factors for stroke** | | | | |
| Atrial fibrillation, n (%) | 8 (8.2) | 4 (10.5) | 4 (6.8) | 0.513 |
| Hypertension, n (%) | 49 (50.5) | 18 (47.4) | 31 (52.5) | 0.619 |
| Diabetes, n (%) | 18 (18.6) | 6 (15.8) | 12 (20.3) | 0.574 |
| Coronary heart disease, n (%) | 16 (16.5) | 8 (21.1) | 8 (13.6) | 0.332 |
| Previous stroke, n (%) | 4 (4.1) | 1 (2.6) | 3 (5.1) | 0.553 |
| Smoking, n (%) | 38 (39.2) | 14 (36.8) | 24 (40.7) | 0.706 |
| Drinking, n (%) | 15 (15.5) | 5 (13.2) | 10 (16.9) | 0.614 |
| **Laboratory findings** | | | | |
| White blood cells, $10^9$/L | 7.20 (5.50, 8.45) | 6.65 (5.35, 8.10) | 7.85 (5.68, 8.53) | 0.299 |
| Monocytes, $10^9$/L | 0.40 (0.30, 0.50) | 0.40 (0.30, 0.50) | 0.40 (0.30, 0.50) | 0.968 |
| Neutrophils, $10^9$/L | 4.60 (3.40, 6.00) | 4.00 (3.00, 5.78) | 5.15 (3.90, 6.13) | 0.594 |
| Eosinophil, $10^9$/L | 0.09 (0.05, 0.20) | 0.11 (0.08, 0.20) | 0.08 (0.04, 0.18) | 0.105 |
| Basophil, $10^9$/L | 0.02 (0.01, 0.03) | 0.02 (0.01, 0.03) | 0.02 (0.01, 0.03) | 0.568 |
| Lymphocytes, $10^9$/L | 1.70 (1.25, 2.00) | 1.80 (1.48, 2.13) | 1.65 (1.20, 2.00) | 0.103 |
| Platelets, $10^9$/L | 227.09 ± 55.65 | 217.82 ± 44.02 | 229.14 ± 67.76 | 0.364 |
| Red blood cells, $10^9$/L | 4.42 ± 0.45 | 4.47 ± 0.46 | 4.33 ± 0.55 | 0.184 |
| Hemoglobin, g/L | 135.50 (126.75, 145.25) | 135.50 (126.00, 148.00) | 136.50 (125.75, 145.25) | 0.520 |
| C-reactive protein, mg/L | 2.40 (1.23, 6.20) | 3.65 (0.83, 4.58) | 1.90 (1.30, 6.72) | 0.951 |
| Fibrinogen, g/L | 3.09 (2.69, 3.59) | 3.07 (2.70, 3.49) | 3.12 (2.65, 3.65) | 0.965 |
| D-dimer, mg/L | 0.49 (0.28, 1.21) | 0.45 (0.28, 1.08) | 0.52 (0.30, 1.27) | 0.426 |
| Cr, μmol/L | 65.00 (53.00, 79.00) | 65.50 (54.75, 81.00) | 62.50 (52.50, 74.00) | 0.740 |
| BUN, mmol/L | 4.94 (4.14, 5.92) | 5.00 (4.51, 6.16) | 4.61 (3.85, 5.52) | 0.989 |
| UA, μmol/L | 289.00 (246.00, 351.50) | 299.00 (249.75, 370.50) | 274.50 (239.25, 337.50) | 0.164 |
| Glucose, mmol/L | 6.50 (5.58, 7.85) | 6.10 (5.01, 7.70) | 6.75 (5.80, 8.30) | 0.154 |
| HbA1c, % | 6.10 (5.88, 6.73) | 6.00 (5.68, 6.63) | 6.15 (5.90, 6.80) | 0.342 |
| Homocysteine, μmol/L | 11.75 (10.00, 14.75) | 12.20 (10.23, 16.53) | 11.00 (9.90, 12.70) | 0.368 |
| GPT, unit/L | 14.00 (11.00, 19.25) | 15.00 (12.00, 21.00) | 14.00 (10.25, 18.00) | 0.104 |
| GOT, unit/L | 20.00 (17.00, 25.00) | 21.00 (18.00, 25.00) | 19.00 (17.00, 23.00) | 0.196 |
| TBil, μmol/L | 13.55 (11.10, 17.85) | 11.90 (9.63, 12.13) | 14.00 (11.73, 17.65) | 0.503 |
| TG, mmol/L | 1.17 (0.86, 1.51) | 1.19 (0.93, 1.57) | 1.14 (0.87, 1.54) | 0.137 |
| TC, mmol/L | 4.48 ± 0.87 | 4.42 ± 0.92 | 4.52 ± 0.84 | 0.548 |
| HDL, mmol/L | 1.27 ± 0.25 | 1.28 ± 0.29 | 1.27 ± 0.23 | 0.823 |
| LDL, mmol/L | 2.53 ± 0.69 | 2.41 ± 0.68 | 2.61 ± 0.69 | 0.159 |
| **PDGF-BB, ng/L** | 495.63 (280.96, 698.06) | 467.50 (275.83, 627.98) | 478.18 (251.66, 689.75) | 0.468 |
| **Stroke onset – to – Blood collection time (h)** | 24.00 (7.25, 48.00) | 33.00 (14.00, 48.00) | 24.00 (7.00, 72.00) | 0.340 |
| **Thrombolysis therapy** | 9 (9.3) | 3 (7.9) | 6 (10.2) | 0.706 |
| **Clinical findings** | | | | |
| NIHSS at admission | 3.00 (2.00, 5.00) | 2.00 (1.00, 3.00) | 5.00 (3.00, 9.00) | < 0.001 |

| | | | | |
|---|---|---|---|---|
| NIHSS at discharge | 2.50 (1.00, 4.00) | 1.00 (0.25, 2.00) | 4.00 (3.00, 8.00) | < 0.001 |

**Note:**

[1]Continuous variables with normal distribution were expressed as mean ± SD and non-normally distributed data were expressed as median (P25, P75). Categorical variables were presented as frequencies.

BUN, blood urea nitrogen; Cr, serum creatinine; GPT, glutamic pyruvic transaminase; GOT, glutamic oxaloacetic transaminase; HbA1c, Hemoglobin A1c; HDL, high density lipoprotein; LDL, low density lipoprotein; NIHSS, National Institute of Health stroke scale; TBil, Total bilirubin; TC, total cholesterol; TG, triglyceride; PDGF-BB, Platelet-derived growth factor-BB; UA, uric acid.

| ID | Description | setSize | enrichmentScore | NES | pvalue | p.adjust | qvalue | rank | leading_edge | core_enrichment | log2err |
|---|---|---|---|---|---|---|---|---|---|---|---|
| HALLMARK_ALLOGRAFT_REJECTION | HALLMARK_ALLOGRAFT_REJECTION | 70 | -0.478118328345098 | -1.95490998830677 | 3.76551383650894e-05 | 0.00184510177988938 | 0.00089377595857497 | 1407 | tags=41%, list=17%, signal=35% | Ube2n/Tap1/Irf8/Tapbp/Icam1/Cd47/Tpd52/Nlrp3/Ifngr2/C2/Stat1/Acvr2a/Abi1/Ripk2/Hcls1/Nme1/Tgfb2/Hif1a/Tgfb1/Ccl19/Prkcg/H2-T23/Capg/Socs1/Jak2/Lyn/Gbp2/Srgn/Stab1 | 0.557332238758646 |
| HALLMARK_TNFA_SIGNALING_VIA_NFKB | HALLMARK_TNFA_SIGNALING_VIA_NFKB | 128 | -0.38879995046861 | -1.78631118055588 | 0.000119488671814004 | 0.00292747245944309 | 0.0014180813937524 | 1231 | tags=27%, list=15%, signal=24% | Fos/Rcan1/Icam1/Ccrl2/Nfe2l2/Ifit2/Ifngr2/Ccn1/Birc2/Nr4a3/Klf2/Irf1/Cebpd/Sqstm1/Fosb/Cflar/Plek/Klf9/Ripk2/Nfkb1/Zbtb10/Maff/Il1a/Nampt/Ppp1r15a/Ets2/Serpine1/Cxcl1/Litaf/Relb/Trib1/Cd83/Egr1/Klf4/Rnf19b | 0.538434096309916 |
| HALLMARK_INTERFERON_ALPHA_RESPONSE | HALLMARK_INTERFERON_ALPHA_RESPONSE | 62 | -0.479509977210494 | -1.91108102482611 | 0.000189522612428665 | 0.00309553600300153 | 0.00149949216273148 | 1842 | tags=42%, list=22%, signal=33% | Nmi/B2m/Parp14/Psme2/Lgals3bp/Parp9/Tap1/Stat2/Trim12c/Cd47/Ccrl2/Rnf31/Ifit2/Irf1/Psma3/Tent5a/Trim26/Ripk2/Cmtr1/Ifitm1/Parp12/Cnp/Elf1/Casp8/Ifitm3/Gbp2 | 0.518848077743792 |
| HALLMARK_INFLAMMATORY_RESPONSE | HALLMARK_INFLAMMATORY_RESPONSE | 82 | -0.42948040289064 | -1.8329933703847 | 0.000372041452642714 | 0.00445279469216594 | 0.00215695463941656 | 1353 | tags=34%, list=16%, signal=29% | Cd14/Slc7a2/Tapbp/Cd82/Icam1/Ccrl2/Dcbld2/Nlrp3/P2rx4/Ifngr2/Ptafr/Irf1/Rgs16/Acvr2a/Kif1b/Abi1/Chst2/Axl/Ripk2/Nfkb1/Il1a/Hif1a/Nampt/Ifitm1/Serpine1/Lyn/Stab1/Csf3r | 0.49849310876659 |
| HALLMARK_INTERFERON_GAMMA_RESPONSE | HALLMARK_INTERFERON_GAMMA_RESPONSE | 106 | -0.385206245821144 | -1.69410267610709 | 0.000454366805323055 | 0.00445279469216594 | 0.00215695463941656 | 1419 | tags=28%, list=17%, signal=24% | Lgals3bp/Tap1/Stat2/Irf8/Tapbp/Icam1/Rnf31/Sppl2a/Ifit2/Nod1/Lysmd2/Irf1/Stat1/Xaf1/Psma3/Pml/Trim26/Ripk2/Mt2/Ptpn2/Nfkb1/Cmtr1/Hif1a/Nampt/Parp12/Casp8/Ifitm3/Socs1/Jak2/Irf5 | 0.49849310876659 |
| HALLMARK_APICAL_JUNCTION | HALLMARK_APICAL_JUNCTION | 102 | 0.35843542804152 | 1.60844594981437 | 0.00411277491991402 | 0.0293911070011418 | 0.0142371901213493 | 1462 | tags=31%, list=18%, signal=26% | Mdk/Ctnna1/Nectin1/Actn4/Pcdh1/Hras/Nrxn2/Stx4a/Pkd1/Ldlrap1/Pbx2/Cdk8/Vcl/Akt3/Myl9/Src/Gnai2/Taok2/Egfr/Nexn/Gtf2f1/Sympk/Jup/Cdh4/Ywhah/Tjp1/Vav2/Icam2/Actn1/Pten/Myh9/Nectin2 | 0.407017918923954 |
| HALLMARK_OXIDATIVE_PHOSPHORYLATION | HALLMARK_OXIDATIVE_PHOSPHORYLATION | 160 | 0.321178168477175 | 1.55022009649812 | 0.00419872957159168 | 0.0293911070011418 | 0.0142371901213493 | 1599 | tags=28%, list=19%, signal=23% | Uqcr11/Etfb/Cox8a/Uqcrq/Uqcr10/Gpx4/Ndufa6/Timm13/Ndufa8/Ndufa2/Ndufb2/Bckdha/Ech1/Cox4i1/Cox5b/Ndufs7/Ndufb7/Ndufs6/Uqcrc1/Ndufv1/Mfn2/Atp6v1f/Cox6c/Atp6v1g1/Alas1/Gpi1/Cpt1a/Atp6v1d/Ndufab1/Polr2f/Idh2/Timm8b/Sucla2/Acaa1a/Vdac3/Cox7c/Glud1/Cyb5r3/Ndufb4/Ndufc1/Slc25a5/Timm9/Cox17/Tcirg1 | 0.407017918923954 |
| HALLMARK_XENOBIOTIC_METABOLISM | HALLMARK_XENOBIOTIC_METABOLISM | 97 | -0.343583878749258 | -1.51290270121078 | 0.0141474808370417 | 0.0866533201268802 | 0.0419752748082815 | 1460 | tags=29%, list=18%, signal=24% | Aldh9a1/Tmbim6/Maoa/Dhrs1/Tnfrsf1a/Acp1/Irf8/Tmem97/Acox3/Crot/Gart/Elovl5/Cat/Sertad1/Mt2/Tgfb2/Comt/Ptges/Acox1/Acp2/Ephx1/Car2/Ets2/Serpine1/Lpin2/Tpst1/Fah/Vtn | 0.380730400722792 |
| HALLMARK_REACTIVE_OXYGEN_SPECIES_PATHWAY | HALLMARK_REACTIVE_OXYGEN_SPECIES_PATHWAY | 36 | 0.441331056331659 | 1.62107810565943 | 0.0196895116073523 | 0.0877078244327512 | 0.0424860816413104 | 1537 | tags=33%, list=19%, signal=27% | Gpx4/Ndufa6/Hhex/Cdkn2d/Sbno2/Ftl1/Prdx2/Ptpa/Sod1/Fes/Ndufb4/Egln2 | 0.352487857583619 |
| HALLMARK_IL6_JAK_STAT3_SIGNALING | HALLMARK_IL6_JAK_STAT3_SIGNALING | 47 | -0.417901477082519 | -1.57503301506402 | 0.0188925248389911 | 0.0877078244327512 | 0.0424860816413104 | 1354 | tags=30%, list=16%, signal=25% | Stat2/Cd14/Tnfrsf21/Tnfrsf1a/Ifngr2/Irf1/Stat1/Cbl/Lepr/Ptpn2/Tgfb1/Cxcl1/Socs1/Csf3r | 0.352487857583619 |
| HALLMARK_UV_RESPONSE_DN | HALLMARK_UV_RESPONSE_DN | 109 | -0.319925282489678 | -1.39808421085736 | 0.0189282384965317 | 0.0877078244327512 | 0.0424860816413104 | 1031 | tags=21%, list=12%, signal=19% | Notch2/Ltbp1/Prkar2b/Ccn1/Nr3c1/Acvr2a/Pik3cd/Atrn/Mgmt/Aggf1/Nfkb1/Lamc1/Zmiz1/Ddah1/Prkca/Tfpi/Atxn1/Col3a1/Serpine1/Nr1d2/Prkce/Dbp/Tgfbr2 | 0.352487857583619 |
| HALLMARK_ESTROGEN_RESPONSE_EARLY | HALLMARK_ESTROGEN_RESPONSE_EARLY | 90 | -0.310217580653666 | -1.33605454902902 | 0.0558558558558559 | 0.195495495495495 | 0.0946989351958952 | 1760 | tags=33%, list=21%, signal=27% | Tubb2b/Tmem164/Rhobtb3/Fkbp4/Farp1/Add3/Slc7a2/Fos/Amfr/Abhd2/Dynlt3/Nrip1/Elovl5/Akap1/Prss23/Xbp1/Rbbp8/Sh3bp5/Bcl2/Olfml3/Rab31/Ptges/Chpt1/Retreg1/Thsd4/Elf1/Jak2/Gab2/Slc1a4/Klf4 | 0.257206466468838 |
| HALLMARK_MTORC1_SIGNALING | HALLMARK_MTORC1_SIGNALING | 129 | -0.287883673638223 | -1.32346933193792 | 0.048671456905081 | 0.195495495495495 | 0.0946989351958952 | 1212 | tags=21%, list=15%, signal=18% | Gmps/Tmem97/Acsl3/Hspa4/Ddit3/Cacybp/Elovl5/Xbp1/Psma3/Sqstm1/Got1/Ppia/Calr/Ube2d3/Tpi1/Pgm1/Ifrd1/Rab1a/Actr3/Pitpnb/Nampt/Ppp1r15a/Coro1a/Aldoa/Slc1a4/Psmd13/Sytl2 | 0.321775918075361 |
| HALLMARK_IL2_STAT5_SIGNALING | HALLMARK_IL2_STAT5_SIGNALING | 106 | -0.300223356992676 | -1.32035551870906 | 0.054249547920434 | 0.195495495495495 | 0.0946989351958952 | 965 | tags=23%, list=12%, signal=20% | Igf2r/P2rx4/Plin2/Rgs16/Xbp1/Spry4/Snx14/Bcl2/Lclat1/Serpinb6a/Prkch/Maff/Nop2/Pdcd2l/Rnh1/Car2/Hipk2/Cyfip1/Capg/Ifitm3/Socs1/Fah/Swap70/Cd83 | 0.261663521711573 |
| HALLMARK_HEDGEHOG_SIGNALING | HALLMARK_HEDGEHOG_SIGNALING | 17 | -0.49449317646241 | -1.46522720897378 | 0.0666666666666667 | 0.217777777777778 | 0.105492577271986 | 808 | tags=35%, list=10%, signal=32% | Pml/Tle1/Cdk6/Ets2/Adgrg1/Ophn1 | 0.245041785430996 |
| HALLMARK_TGF_BETA_SIGNALING | HALLMARK_TGF_BETA_SIGNALING | 46 | -0.342044848608479 | -1.28282049731095 | 0.105357142857143 | 0.32265625 | 0.156296201259556 | 936 | tags=26%, list=11%, signal=23% | Ifngr2/Ppm1a/Ube2d3/Smad1/Hdac1/Wwtr1/Rab31/Fkbp1a/Tgfb1/Ppp1r15a/Hipk2/Serpine1 | 0.183023938384487 |
| HALLMARK_GLYCOLYSIS | HALLMARK_GLYCOLYSIS | 105 | -0.269194996261603 | -1.18049683950614 | 0.156190476190476 | 0.450196078431373 | 0.218077092511837 | 1650 | tags=26%, list=20%, signal=21% | Ecd/Fkbp4/Idua/B4galt4/Alg1/Stmn1/Aldh9a1/Pmm2/Mdh2/Gmppa/Sdc3/Pgam1/Got1/Ppia/Fut8/Kif2a/Chst2/Tpi1/Ext1/Ppp2cb/Sdhc/Plod1/Kdelr3/Tpst1/Agrn/Aldoa/Mertk | 0.153158808683073 |
| HALLMARK_KRAS_SIGNALING_UP | HALLMARK_KRAS_SIGNALING_UP | 87 | -0.270479803867208 | -1.16924608510977 | 0.171428571428571 | 0.466666666666667 | 0.226055522725685 | 1647 | tags=28%, list=20%, signal=22% | Evi5/Tor1aip2/Zfp639/Gng11/Gypc/Fcer1g/Irf8/Sdccag8/Laptm5/Crot/Prrx1/Dcbld2/Rgs16/Cbl/Mtmr10/Dnmbp/Ppp1r15a/Tfpi/Car2/Cab39l/Ano1/Trib1/Klf4/Adgrl4 | 0.14551614609036 |
| HALLMARK_ANGIOGENESIS | HALLMARK_ANGIOGENESIS | 20 | -0.40563818905948 | -1.23272865879042 | 0.212371134020619 | 0.520309278350515 | 0.252040255492607 | 627 | tags=25%, list=8%, signal=23% | Thbd/Fstl1/Col3a1/Vtn/Kcnj8 | 0.135002033828631 |
| HALLMARK_MITOTIC_SPINDLE | HALLMARK_MITOTIC_SPINDLE | 141 | 0.240908192507071 | 1.1275133983631 | 0.210526315789474 | 0.520309278350515 | 0.252040255492607 | 1612 | tags=24%, list=20%, signal=20% | Clip2/Pcnt/Actn4/Rasal2/Nck1/Itsn1/Rock1/Dynll2/Wasf2/Rictor/Vcl/Cdc42ep4/Abr/Sptbn1/Nin/Taok2/Plekhg2/Arl8a/Arhgap27/Dync1h1/Ccdc88a/Nedd9/Map3k11/Cttn/Ralbp1/Pcm1/Klc1/Smc4/Myh9/Cdc42bpa/Pkd2/Pafah1b1/Arfgef1/Csnk1d | 0.137250779610133 |
| HALLMARK_WNT_BETA_CATENIN_SIGNALING | HALLMARK_WNT_BETA_CATENIN_SIGNALING | 32 | 0.325442918667669 | 1.13533856547609 | 0.269461077844311 | 0.523093681917211 | 0.253389033643497 | 730 | tags=19%, list=9%, signal=17% | Hdac5/Ncor2/Jag2/Ccnd2/Adam17/Notch1 | 0.115734447701689 |
| HALLMARK_UNFOLDED_PROTEIN_RESPONSE | HALLMARK_UNFOLDED_PROTEIN_RESPONSE | 86 | -0.262566338346294 | -1.11287499615703 | 0.28219696969697 | 0.523093681917211 | 0.253389033643497 | 2564 | tags=38%, list=31%, signal=27% | Lsm1/Banf1/Serp1/Ywhaz/Paip1/Npm1/Slc7a5/Wfs1/Nolc1/Mtrex/Eif4a1/Arfgap1/Atf3/Nfya/Sec31a/Sec11a/Atp6v0d1/Atf6/Parn/Herpud1/Shc1/Nop56/Atf4/Xbp1/Calr/Pop4/Exosc4/Nfyb/Spcs1/Kdelr3/Eif4e/Exosc2/Slc1a4 | 0.109250039480711 |
| HALLMARK_ESTROGEN_RESPONSE_LATE | HALLMARK_ESTROGEN_RESPONSE_LATE | 88 | -0.254493599833281 | -1.10623759426424 | 0.247723132969035 | 0.523093681917211 | 0.253389033643497 | 1297 | tags=26%, list=16%, signal=22% | Mocs2/Fos/Amfr/Hspa4l/Abhd2/Dynlt3/Nrip1/Prkar2b/Elovl5/Prss23/Xbp1/Rbbp8/Bcl2/Fgfr3/Rab31/Ptges/Jak1/Chpt1/Gper1/Car2/Jak2/Slc1a4/Klf4 | 0.115240003147781 |
| HALLMARK_G2M_CHECKPOINT | HALLMARK_G2M_CHECKPOINT | 104 | -0.24795143594778 | -1.1005632201313 | 0.261029411764706 | 0.523093681917211 | 0.253389033643497 | 2231 | tags=33%, list=27%, signal=24% | Rad21/Kmt5a/Prmt5/Nolc1/Nup98/Ilf3/Tle3/Gspt1/Odf2/Tra2b/Hspa8/Stmn1/Abl1/Smad3/Bub3/Mcm6/Dr1/Odc1/Orc5/Cks2/Ncl/Pttg1/Smc1a/Notch2/Ccnt1/Chmp1a/Smarcc1/Pml/Mt2/G3bp1/Hif1a/Rad23b/Tgfb1/Cdk4 | 0.112378519110475 |
| HALLMARK_P53_PATHWAY | HALLMARK_P53_PATHWAY | 126 | 0.234653924959608 | 1.09029466771889 | 0.269565217391304 | 0.523093681917211 | 0.253389033643497 | 1362 | tags=23%, list=17%, signal=20% | Hint1/Mxd4/Hras/Btg1/Cgrrf1/Gadd45a/H2aj/Jag2/Tsc22d1/Ccnd2/Fbxw7/Nupr1/Plxnb2/Pdgfa/Rrad/Xpc/Notch1/Blcap/Tm7sf3/Vwa5a/Rb1/Prmt2/Dnttip2/Cyfip2/Tob1/Trafd1/Sec61a1/Prkab1/Rab40c | 0.121543282035469 |

| | | | | | | | | | | | |
|---|---|---|---|---|---|---|---|---|---|---|---|
| HALLMARK_DNA_REPAIR | HALLMARK_DNA_REPAIR | 105 | 0.24078703699751 | 1.07439273882557 | 0.276729559748428 | 0.523093681917211 | 0.253389033643497 | 2037 | tags=29%, list=25%, signal=22% | Bola2/Gpx4/Bcam/Eif1b/Pold3/Xpc/Rala/Gtf2f1/Itpa/Polr2e/Polr2f/Rev3l/Srsf6/Pde4b/Sec61a1/Mpc2/Polr2c/Pold4/Cox17/Polb/Arl6ip1/Rpa2/Ercc1/Adrm1/Edf1/Polr2i/Nt5c/Pole4/Nfx1/Supt4a | 0.117249716749792 |
| HALLMARK_FATTY_ACID_METABOLISM | HALLMARK_FATTY_ACID_METABOLISM | 97 | -0.243435087304402 | -1.07191758383136 | 0.288235294117647 | 0.523093681917211 | 0.253389033643497 | 2218 | tags=33%, list=27%, signal=24% | Kmt5a/Pdha1/Pdhb/Adsl/Etfdh/Suclg2/Fh1/Cpox/Hsd17b10/Gpd2/Serinc1/Hsp90aa1/Idh3g/Aldh9a1/Maoa/Decr1/Odc1/Idh3b/Mdh2/Eci2/Elovl5/Suclg1/Rap1gds1/Car4/Sdhc/Urod/Acsl5/Ptprg/Acox1/Ephx1/Car2/Aldoa | 0.110122263835536 |
| HALLMARK_MYC_TARGETS_V2 | HALLMARK_MYC_TARGETS_V2 | 31 | -0.307014783572243 | -1.05325106921002 | 0.375745526838966 | 0.634880372934805 | 0.307539031225833 | 2324 | tags=42%, list=28%, signal=30% | Npm1/Ndufaf4/Nolc1/Hspd1/Mphosph10/Pprc1/Imp4/Tmem97/Nop56/Gnl3/Las1l/Nop2/Cdk4 | 0.0946646223153342 |
| HALLMARK_COMPLEMENT | HALLMARK_COMPLEMENT | 105 | 0.229745045155093 | 1.02512332629574 | 0.371069182389937 | 0.634880372934805 | 0.307539031225833 | 2189 | tags=35%, list=27%, signal=26% | Calm1/Usp14/Stx4a/Gnb2/Csrp1/Cp/Irf7/Ctsh/Ctsb/Src/Gnai2/Dgkh/Raf1/Fn1/Lta4h/Kcnip3/Lrp1/Ctsc/Pik3ca/Vcpip1/Pik3r5/Ehd1/C1qc/Dgkg/Dock9/Xpnpep1/Rhog/Spock2/Zeb1/Gng2/Mmp14/Cfh/Cebpb/Cblb/Atox1/Ctsd/Clu | 0.0985499803531691 |
| HALLMARK_MYC_TARGETS_V1 | HALLMARK_MYC_TARGETS_V1 | 158 | -0.219202055855029 | -1.02741601643113 | 0.391941391941392 | 0.64017094017094 | 0.310101806816004 | 2660 | tags=39%, list=32%, signal=27% | Hdac2/Eif1ax/Prdx3/Hnrnpu/Canx/Ncbp2/Syncrip/Pold2/Psma4/Cops5/Vdac1/Erh/Cct2/Psmc4/Npm1/Tufm/Eef1b2/Nolc1/Pcbp1/Tomm70a/Eif4a1/Hspd1/Tcp1/Ssb/Ywhaq/Gspt1/Ilf2/Psmc6/Tra2b/Prpf31/Fbl/Phb2/Bub3/Ruvbl2/Hnrnpr/Mcm6/Odc1/Eif4h/Srsf7/Acp1/Srsf3/Ddx21/Psmb3/C1qbp/Hddc2/Snrpg/Nop56/Smarcc1/Ppia/Gnl3/Hsp90ab1/Rrm1/Ifrd1/Psma6/Set/Nme1/G3bp1/Eif4e/Rad23b/Stard7/Slc25a3/Cdk4 | 0.0875697115548511 |
| HALLMARK_COAGULATION | HALLMARK_COAGULATION | 59 | -0.259360926007472 | -1.01741078647876 | 0.42970297029703 | 0.658685150178534 | 0.319070177004118 | 2642 | tags=46%, list=32%, signal=31% | Mmp11/C1s1/Casp9/Lamp2/Lgmn/Wdr1/Klf7/Adam9/Bmp1/Ctso/Plat/Gsn/Htra1/Pdgfb/Cpq/Iscu/Vwf/C2/Prss23/Plek/Thbd/Rac1/Sparc/Maff/Capn2/Anxa1/Serpine1 | 0.086794975677172 |
| HALLMARK_ADIPOGENESIS | HALLMARK_ADIPOGENESIS | 154 | 0.210759448634858 | 1.00558070149539 | 0.438297872340426 | 0.658685150178534 | 0.319070177004118 | 367 | tags=9%, list=4%, signal=9% | Uqcr11/Etfb/Cox8a/Uqcrq/Uqcr10/Gpx4/Ifngr1/Bckdha/Ech1/Ucp2/Gadd45a/Itsn1/Ndufb7/Uqcrc1 | 0.0897104652742981 |
| HALLMARK_PI3K_AKT_MTOR_SIGNALING | HALLMARK_PI3K_AKT_MTOR_SIGNALING | 69 | -0.244583403878966 | -0.998327971113006 | 0.458089668615984 | 0.658685150178534 | 0.319070177004118 | 989 | tags=19%, list=12%, signal=17% | Ddit3/Nod1/Sqstm1/Calr/Arpc3/Ube2d3/Rac1/Actr3/Eif4e/Mapkap1/Smad2/Cab39l/Cdk4 | 0.0824344091527096 |
| HALLMARK_HEME_METABOLISM | HALLMARK_HEME_METABOLISM | 126 | -0.220422774760663 | -0.997650946919896 | 0.483394833948339 | 0.658685150178534 | 0.319070177004118 | 1721 | tags=26%, list=21%, signal=21% | Daam1/Clcn3/Synj1/Bnip3l/Ppp2r5b/Fech/Gypc/Marchf2/Kdm7a/Htra2/Dcaf11/Riok3/Gmps/Ubac1/Tmem9b/Mkrn1/Nr3c1/Mocos/Cat/Cir1/Adipor1/Isca1/Lrp10/Slc30a1/Minpp1/Urod/Mboat2/Bmp2k/Ypel5/Bach1/Car2/Lpin2/Lmo2 | 0.0766746872490091 |
| HALLMARK_HYPOXIA | HALLMARK_HYPOXIA | 128 | -0.215825037595857 | -0.991591375556259 | 0.483931947069943 | 0.658685150178534 | 0.319070177004118 | 989 | tags=17%, list=12%, signal=15% | Ddit3/Tpd52/Ccn1/Plin2/Sdc3/Ampd3/Nr3c1/Bcl2/Chst2/Tpi1/Ext1/Pgm1/Kdm3a/Mt2/Maff/Kdelr3/Prkca/Ppp1r15a/Errfi1/Serpine1/Aldoa/Tgfb3 | 0.0778840111977068 |
| HALLMARK_BILE_ACID_METABOLISM | HALLMARK_BILE_ACID_METABOLISM | 49 | -0.260524271177534 | -0.97961886388575 | 0.48314606741573 | 0.658685150178534 | 0.319070177004118 | 1724 | tags=24%, list=21%, signal=19% | Abcd3/Pex7/Aldh9a1/Crot/Lipe/Scp2/Gstk1/Abca5/Cat/Abca8b/Acsl5/Atxn1 | 0.0774767547848578 |
| HALLMARK_KRAS_SIGNALING_DN | HALLMARK_KRAS_SIGNALING_DN | 38 | -0.269091401827556 | -0.959351641142874 | 0.536679536679537 | 0.709802028005794 | 0.343831432441288 | 1620 | tags=29%, list=20%, signal=23% | Idua/Gtf3c5/Slc25a23/Asb7/Mast3/Kmt2d/Fgfr3/Chst2/Tgfb2/Kcnmb1/Nr6a1 | 0.0736212741014394 |
| HALLMARK_UV_RESPONSE_UP | HALLMARK_UV_RESPONSE_UP | 102 | -0.215782576988094 | -0.942822241858919 | 0.55045871559633 | 0.709802028005794 | 0.343831432441288 | 2211 | tags=34%, list=27%, signal=25% | Dlg4/Wiz/Tfrc/Hspa13/Polg2/Cyb5r1/Bak1/Bcl2l11/Nxf1/Atf3/Stk25/Stard3/Fkbp4/Parp2/Tmbim6/Maoa/Tap1/Fos/Cdc5l/Icam1/Ddx21/Dnaja1/E2f5/Irf1/Sqstm1/Fosb/Tgfbrap1/Urod/Ephx1/Eif5/Cnp/Car2/Cxcl1/Aldoa/Lyn | 0.0697792517076235 |
| HALLMARK_APICAL_SURFACE | HALLMARK_APICAL_SURFACE | 18 | -0.307407429729152 | -0.91686738922683 | 0.570610687022901 | 0.716921119592875 | 0.34727995380568 | 3059 | tags=61%, list=37%, signal=39% | App/Adipor2/Shroom2/Hspb1/Rtn4rl1/Sulf2/Scube1/Dcbld2/Atp8b1/Ncoa6/Lyn | 0.0699458736501659 |
| HALLMARK_MYOGENESIS | HALLMARK_MYOGENESIS | 97 | -0.211012788944693 | -0.929152495590236 | 0.590196078431373 | 0.722990196078431 | 0.35021984292869 | 584 | tags=13%, list=7%, signal=13% | Ifrd1/Wwtr1/Sparc/Sptan1/Ptp4a3/Mapk12/Sod3/Casq2/Tgfb1/Sorbs3/Cnn3/Col3a1/Agrn | 0.0696133438850965 |
| HALLMARK_APOPTOSIS | HALLMARK_APOPTOSIS | 96 | -0.21382858274886 | -0.921468395731612 | 0.626334519572954 | 0.748546133148164 | 0.362599258742354 | 1810 | tags=28%, list=22%, signal=22% | Atf3/Plat/Gsn/Cyld/Bnip3l/Ppp2r5b/Tgfbr3/Ptk2/Tap1/Cd14/Dnaja1/Ddit3/Igf2r/Irf1/Sqstm1/Diablo/Cflar/Mgmt/Sptan1/Vdac2/Tgfb2/Il1a/Anxa1/Gstm2/Rhot2/Casp8/Ifitm3 | 0.0619762736001998 |
| HALLMARK_EPITHELIAL_MESENCHYMAL_TRANSITION | HALLMARK_EPITHELIAL_MESENCHYMAL_TRANSITION | 102 | -0.197054296463439 | -0.860992468218797 | 0.772477064220183 | 0.880841223439068 | 0.426683621148966 | 1031 | tags=17%, list=12%, signal=15% | Notch2/Calu/Ccn1/Eln/Loxl2/Fstl1/Sparc/Col4a1/Plod1/Lamc1/Colgalt1/Tgfb1/Ecm2/Capg/Col3a1/Serpine1/Cxcl1 | 0.0535669384770951 |
| HALLMARK_SPERMATOGENESIS | HALLMARK_SPERMATOGENESIS | 44 | -0.217509276717808 | -0.821548865777383 | 0.772983114446529 | 0.880841223439068 | 0.426683621148966 | 2252 | tags=32%, list=27%, signal=23% | Chfr/Pias2/Gpr182/Rad17/Lpin1/Clpb/Agfg1/Parp2/Hspa4l/Pebp1/Ip6k1/Topbp1/Art3/Septin4 | 0.0545680646670941 |
| HALLMARK_PEROXISOME | HALLMARK_PEROXISOME | 67 | -0.19998051836464 | -0.820873321070905 | 0.817142857142857 | 0.91 | 0.440808269315086 | 1724 | tags=25%, list=21%, signal=20% | Abcd3/Prdx1/Slc25a17/Cadm1/Vps4b/Aldh9a1/Cln8/Eci2/Pex11b/Scp2/Elovl5/Gstk1/Smarcc1/Cat/Acsl5/Acox1/Atxn1 | 0.0526973085708419 |
| HALLMARK_PROTEIN_SECRETION | HALLMARK_PROTEIN_SECRETION | 85 | -0.185183628740025 | -0.785625254348717 | 0.888461538461538 | 0.920751633986928 | 0.446016411260237 | 1849 | tags=24%, list=22%, signal=18% | Cd63/Arfip1/Anp32e/Clcn3/Sec31a/Vps4b/Ocrl/Cln5/Tmed2/Tpd52/Igf2r/Rab2a/Tsg101/Kif1b/Bnip3/Rab22a/Ap3b1/Copb1/Mon2/Gnas | 0.0492917662312669 |
| HALLMARK_E2F_TARGETS | HALLMARK_E2F_TARGETS | 97 | -0.176061298149291 | -0.775250615711008 | 0.901960784313726 | 0.920751633986928 | 0.446016411260237 | 1807 | tags=22%, list=22%, signal=17% | Gspt1/Tra2b/Ccp110/Anp32e/Eed/Stmn1/Nup205/Mcm6/Ppp1r8/Prkdc/Pnn/Cks2/Pttg1/Smc1a/Nop56/Ipo7/Trp53/Tubb5/Nme1/Exosc8/Cdk4 | 0.0494904866135495 |
| HALLMARK_ANDROGEN_RESPONSE | HALLMARK_ANDROGEN_RESPONSE | 61 | -0.191101584428487 | -0.772553075802287 | 0.887429643527205 | 0.920751633986928 | 0.446016411260237 | 1443 | tags=18%, list=17%, signal=15% | Vapa/Steap4/Adamts1/Acsl3/Abhd2/Tpd52/Elovl5/Zbtb10/Zmiz1/Cdk6/Mertk | 0.0482149710025564 |
| HALLMARK_NOTCH_SIGNALING | HALLMARK_NOTCH_SIGNALING | 23 | -0.220063595533672 | -0.689993963983576 | 0.894538606403013 | 0.920751633986928 | 0.446016411260237 | 2419 | tags=43%, list=29%, signal=31% | Maml2/Notch3/Hes1/Dtx4/Jag1/Dtx2/Heyl/Notch2/Tcf7l2/Prkca | 0.0480220415968251 |
| HALLMARK_CHOLESTEROL_HOMEOSTASIS | HALLMARK_CHOLESTEROL_HOMEOSTASIS | 42 | -0.173125643808158 | -0.639047351785873 | 0.977596741344195 | 0.977596741344195 | 0.473552447956047 | 324 | tags=7%, list=4%, signal=7% | S100a11/Atxn2/Errfi1 | 0.0475434194201301 |
